# The PhageExpressionAtlas reveals shared and unique transcriptional patterns across phage-host interactions

**DOI:** 10.64898/2026.03.30.715471

**Authors:** Maik Wolfram-Schauerte, Caroline Trust, Nils Waffenschmidt, Kay Nieselt

## Abstract

Time-resolved transcriptomic profiling has been used to study phage-host interactions for more than a decade. However, the resulting datasets are not readily accessible for custom re-analysis, and resources are lacking that provide standardized processing, storage, and analysis of transcriptomes from phage infections. Here, we present the PhageExpressionAtlas, the first bioinformatics resource for storing time-resolved dual RNA-sequencing data from phage infections. This data was processed uniformly using a custom analysis pipeline and is presented for interactive exploration through visualisation. The PhageExpressionAtlas currently hosts 42 datasets from 23 studies. Using the PhageExpressionAtlas, we replicate key findings from original publications and extend hypothesis testing across multiple phage-host systems. By systematically querying and analyzing the underlying database, we evaluate approaches to phage gene classification and show that uncharacterized phage genes are expressed across all infection phases. Moreover, we provide a comprehensive view of the expression dynamics of anti-phage defenses as well as host- and phage-encoded anti-defense systems in the infection context, indicating unique and conserved patterns of transcriptional regulation underlying bacterial anti-phage immunity and phage counter-strategies. Together, the PhageExpressionAtlas is a unifying resource that democratizes transcriptomics-driven analyses of phage-host interactions and supports integrative cross-study assessment.

## Introduction

Bacteriophages (phages) are viruses that infect bacteria and can lyse their hosts, thereby shaping microbial communities across diverse ecosystems. In recent years, phages have also gained increasing relevance in biotechnology and medicine. Phage-derived enzymes widely used in molecular biology [1], anti-phage defense systems such as CRISPR-Cas [2, 3], phage-encoded anti-defense strategies that counteract these systems [4], and additional mechanisms enabling host takeover [5, 6] have become central topics of investigation. In parallel, antibiotic resistance continues to rise as a major global health threat, motivating renewed interest in phages as alternatives or complements to antibiotics, including phage therapy [7].

To fully exploit the potential of phages and their genomic repertoires, a mechanistic understanding of phage-host interactions is essential. One key approach is dual RNA sequencing (dual RNA-seq), which simultaneously quantifies phage and host transcriptomes across infection stages [8]. Time-resolved dual RNA-seq studies across diverse phage infections have yielded fundamental insights into host responses, including anti-phage defense, as well as phage anti-defense and takeover programs and their interplay [9, 10, 11, 12, 13]. While these studies illuminate specific facets of individual phage-host systems, their primary analyses are typically framed by the hypotheses and functional annotations available at the time. As phage research continues to uncover new interaction mechanisms and gene functions, previously generated dual RNA-seq datasets become increasingly valuable for re-analysis and cross-study comparison, yet are rarely revisited in a systematic manner.

Notably, dual RNA-seq data are typically not processed in a standardized manner and are rarely made available through dedicated databases in a format that supports rapid re-analysis. Moreover, visual exploration options that are accessible to non-bioinformaticians remain limited, with only few exceptions [10]. As a consequence, comprehensive and systematic analyses - and especially comparisons across phage infections - are still lacking, despite their potential to substantially advance mechanistic understanding and to inform applications in biotechnology and therapy. It is therefore instrumental to provide a resource that enables access to, and interactive visualization and exploration of, transcriptome-resolved phage infections. While databases and tools for phage genomics and phenotypes are available [14, 15], comparable resources that facilitate gene expression-centric analyses are currently missing.

Here, we present the PhageExpressionAtlas, the first resource in phage research that enables interactive, visual exploration of phage-host interactions at the transcriptome level. To this end, we implemented a unified processing and analysis framework and uniformly processed 42 public time-resolved dual RNA-seq datasets spanning diverse phage-host systems across infection stages from 23 studies. The atlas includes multiple clinically relevant hosts, including pathogens such as *Staphylococcus aureus* and *Pseudomonas aeruginosa*, and covers both model and therapeutic phages, thereby representing a broad spectrum of infection modalities. Users can query the collection, download processed data, and visualize phage and host gene expression dynamics across infection phases for selected phage-host interactions using heatmaps and profile plots. In addition, PhageExpressionAtlas supports classification of phage genes into early, middle, and late expression classes, visualization of their temporal profiles, and inspection of their genomic context. This functionality provides insights into infection-stage-specific programs, supports functional annotation of phage genes, and facilitates investigation of genome organization. Leveraging the aggregated dataset collection, we systematically assess the transcriptional dynamics of bacterial anti-phage defense systems and identify programmed expression of phage-encoded anti-defense systems, extending current knowledge by placing defense and counter-defense modules into their native infection contexts.

Altogether, the PhageExpressionAtlas provides a foundation for a growing visualization and analysis resource that democratizes transcriptomics-driven phage research and enables integrative cross-study assessment. The PhageExpressionAtlas is available at https://phageexpressionatlas.cs.uni-tuebingen.de.

## Materials & Methods

Collection and curation of dual RNA-seq datasets of phage infections

### Data collection

We searched PubMed, GEO, and ENA for dual RNA-seq datasets of phage infections. Datasets were included in the PhageExpressionAtlas if they met the following criteria:

- the dual RNA-seq data cover at least two time points during phage infection,
- raw sequencing data are publicly available or, if not publicly available, were provided by the authors upon request,
- sufficient information on the phage and host strains, including an appropriate reference genome, is available.

Supplementary Table S1A summarizes all data accessions, associated study metadata, and the reference genome accessions used, which were retrieved from the respective repositories. Overall, the collection spans a diverse range of phages - both temperate and virulent - and hosts (Fig. 1A,B).

**Fig. 1.**
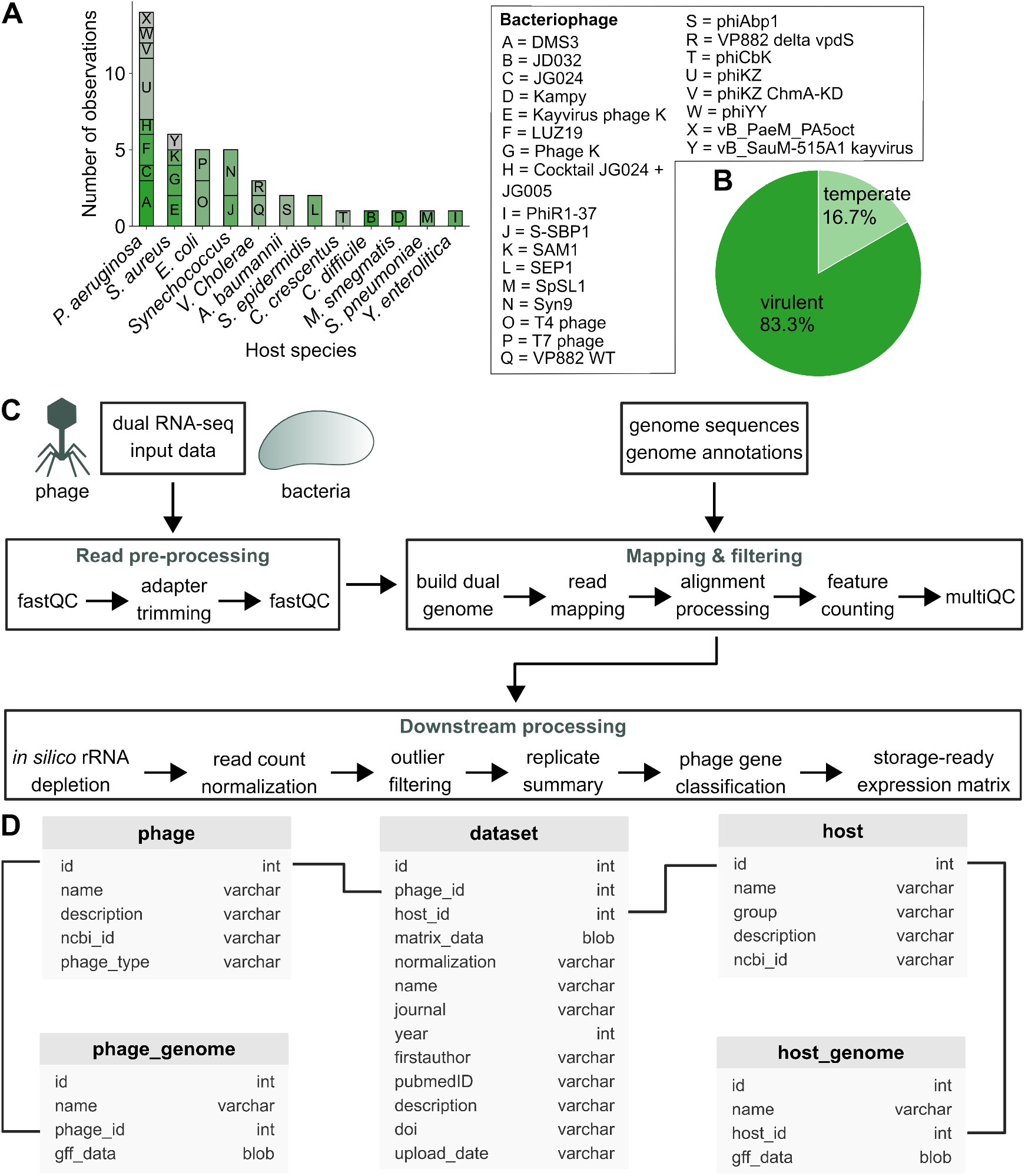
Features, processing and storage of dual RNA-seq datasets in the PhageExpressionAtlas. **A**) Overview of datasets collected for and represented in the PhageExpressionAtlas grouped by host species (summarizing different strains of the same species) and the phages infecting the hosts. **B**) Pie chart illustrating the fractions of temperate and virulent phages across all datasets. **C**) Schematic illustration of key steps of dual RNA-seq data processing using the custom Nextflow pipeline and interactive downstream processing. Processing requires host and phage genome information alongside the raw dual RNA-seq data. **D**) Schematic representation of the SQlite database, in which processed dual RNA-seq data are stored (“matrix data’) in four different formats alongside phage and host information and the corresponding genome annotations.

### Data processing

For unified processing of raw RNA-seq reads, we implemented a Nextflow [16] pipeline that performs all computational steps required to generate gene-level count tables (Fig. 1C). The pipeline comprises quality control with FastQC (v0.12.1) [17], adapter trimming with Cutadapt (v4.9) [18], and read mapping with HISAT2 [19] against a combined reference consisting of the host and phage genomes. Alignments were processed with Samtools (v1.6), and gene-level quantification was performed with featureCounts (v2.0.1). We retained primary alignments only, while allowing multi-mapping reads and reads overlapping multiple features to be counted. Finally, we employed MultiQC (v1.32) to summarize quality-control and processing metrics across samples.

To further process the resulting count tables, we implemented a Jupyter Notebook that applies a consistent post-processing workflow across datasets while permitting dataset-specific adaptations where required (Fig. 1C). Count tables were depleted of rRNA features retrieved from the genome annotation, followed by per-sample transcripts-per-million (TPM) normalization:

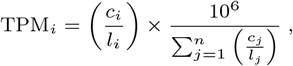

where the TPM of gene *i* in a given sample is computed from its read count *c*_*i*_ and gene length *l*_*i*_, and normalized by the sum of length-normalized counts across all *n* genes in that sample.

Replicate consistency was assessed by principal component analysis (PCA), and outlier replicates were removed in rare cases. Replicates were then aggregated within each time point by computing the mean of TPM values (“TPM means”), which serves as the basis for the heatmaps displayed in the PhageExpressionAtlas. Importantly, we calculated a mean-stabilized variance by dividing the variance of each gene’s expression over time by its mean expression over time, which we use to rank order genes to display gene subsets in heatmaps.

To visualize temporal expression profiles, we further computed “fractional” expression from TPM means by dividing each gene’s TPM value at each time point by its maximal TPM value across all time points.

Phage gene classification assigns phage genes to early, middle, or late infection phases based on their temporal expression patterns. Following common practice in the field, we pre-computed dataset-specific expression classes during processing using study-derived phase boundaries on replicate aggregated TPM values (TPM means). Specifically, for each gene we determined whether its expression exceeded a fraction *θ* of its maximal expression across time points and then assigned the class according to the corresponding time bin:

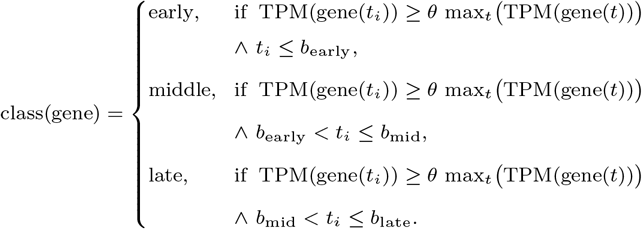

where *θ* is a relative expression threshold and *b*_early_, *b*_mid_, and *b*_late_ denote dataset-specific boundaries for the early, middle, and late infection phases, respectively. For pre-computed classes, we used *θ* = 1 (“Class Max”) and *θ* = 0.2 (“Class Threshold”). In most datasets, these settings recapitulated the phase-resolved gene class distributions reported in the original studies.

In addition, the atlas supports user-defined classification based on custom infection-phase boundaries (*b*_early_, *b*_mid_, *b*_late_) and a user-selected threshold *θ* relative to each gene’s maximal expression across time points. In addition to TPM, we generated a log_2_+1-transformed representation of the data.

To enable efficient storage and querying of host and phage genomes, GFF files were restricted to `gene` features and reformatted. Where overlapping gene symbols occurred, we generated unique symbols; if no symbol was available, the gene identifier was used. We further improved the annotation of protein-coding phage genes by running Pharokka (v1.8.2) [20] with default settings on the protein FASTA files sorted by genomic coordinates, and we added resulting annotations and functional categories to the re-formatted GFF files. For annotated non-coding features (e.g., tRNAs), the corresponding biotype (tRNA, ncRNA, etc.) was added to the category field.

### Database creation

We implemented an SQLite3 database to store all data underlying the PhageExpressionAtlas (Fig. 1D). The database comprises five tables. The central table (“dataset”) stores dataset-level records together with associated study metadata and identifiers for the corresponding phage and host. For each dataset, we store four expression matrices to support different downstream analyses and visualizations: (i) the raw count matrix, (ii) the per-sample TPM-normalized matrix, (iii) mean TPM values aggregated per time point (“TPM means”), and (iv) the matrix of fractional expression values. Each matrix is stored as a pickled `pandas` DataFrame. Phage and host identifiers link to dedicated tables containing organism-level metadata, which in turn link to genome annotations derived from the corresponding GFF files and likewise stored as pickled `pandas` DataFrames.

We provide the database creation workflow as a Jupyter Notebook, which can be adapted to build a custom database and to run a local instance of the PhageExpressionAtlas. This enables private deployment, for example, to explore datasets that cannot yet be shared publicly.

### PhageExpressionAtlas frontend and backend

#### Interface design and functionalities

Building on this foundation, we developed the PhageExpressionAtlas as an interactive web application to facilitate transcriptomics-based (re-)investigation of phage infections. The atlas enables users to search the database and visually explore selected datasets representing phage-host interactions of interest using common visualizations such as heatmaps and profile plots. In addition, expression-based classification of phage genes into infection phases supports hypothesis generation and functional interpretation of phage genes.

We structured the PhageExpressionAtlas into five pages. The landing page (“Home”) introduces the atlas, summarizes available exploration options of the website, and provides a brief background on the relevance of phage research and the role of dual transcriptomics for this (Supplementary Fig. S2).

The “Data Overview” page summarizes dataset characteristics as well as processing and storage in the database of the PhageExpressionAtlas (Fig. 1C,D; Supplementary Fig. S3). It also provides descriptive statistics on the distribution of hosts and phages across included datasets (Supplementary Fig. S3A-C). A comprehensive, searchable table allows users to retrieve dataset details, including links to the original publications (Supplementary Fig. S3D). For each dataset, dedicated buttons link to the “Dataset Exploration” or “Genome Viewer” pages with the corresponding dataset pre-selected.

The “Dataset Exploration” page serves as the core interactive visualization interface of the PhageExpressionAtlas (Supplementary Fig. S4,5). Users can select a phage, host, and dataset of interest and obtain an overview of time-resolved expression dynamics (Supplementary Fig. S4A). In total, three visualization panels are displayed. The upper panel displays a heatmap of z-score-normalized expression (per gene) for all phage genes (left) based on mean TPM values per time point (Supplementary Fig. S4B). On the right, a corresponding heatmap shows the same number of the most variable host genes. As for all visualizations in the atlas, help buttons provide contextual information about the displayed plots, the underlying data, and available selection options.

The middle panel provides the results of phage gene classification functionality. Fractional expression profiles across the measured time points are shown for each phage gene as profile plots and colored according to user-defined classification criteria (Supplementary Fig. S4C). Users can select pre-computed classes (“Class Max” and “Class Threshold”) or perform custom classification by defining infection-phase boundaries via time-point thresholds and a relative expression threshold with respect to each gene’s maximal expression over the infection time course. The profile plots provide immediate visual feedback on the resulting classification, enabling rapid qualitative assessment given the observed expression trajectories.

The lower panel enables generation of z-score-normalized heatmaps and profile plots for user-selected phage and host genes (Supplementary Fig. S5). In addition, for phage genes, also a linear genome track is shown above the plots, allowing genes to be selected in their genomic context and highlighted to indicate the current selection. This track is also linked to the “Genome Viewer” page. This facilitates exploration of specific genomic loci and their transcriptional dynamics. An analogous selection workflow is available for host genes, but without an integrated genome visualization (Supplementary Fig. S5).

All plots on the “Dataset Exploration” page can be downloaded, and phage gene classes generated in the classification panel can be exported. Beyond these dedicated components, interactivity is supported throughout the page via linked selections.

The “Genome Viewer” page provides an additional perspective on time-resolved phage transcriptomes by integrating transcriptomic and genomic information (Supplementary Fig. S6). Users can select a phage and the corresponding dataset, after which phage genes are classified using one of the three options also available on the “Dataset Exploration” page. The phage genome is then displayed as a circular genome plot, with an inner ring indicating functional categories (e.g., DNA, RNA, and nucleotide metabolism) and an outer ring highlighting the inferred temporal class (early, middle, late). Each ring consists of two tracks indicating the forward and reverse strand.

A dynamically adjustable selection window allows users to zoom into a genomic locus of interest and display it as a linear view on the right-hand side (Supplementary Fig. S6). In this view, individual genes and their functional annotations and temporal classes can be interactively inspected, facilitating interpretation of transcriptional programs in their genomic context. For example, the T4 genome illustrates the organization of genes into loci with temporally coordinated transcription (Supplementary Fig. S6), consistent with prior work [21, 10, 22].

To support navigation of the PhageExpressionAtlas, we provide a comprehensive “Help” page that includes written instructions and video demonstrations for each page of the atlas.

#### Implementation

The PhageExpressionAtlas frontend was implemented in HTML, CSS, and JavaScript (JS). For UI elements (e.g., components and buttons), we used the open-source web component library Shoelace (v2.20.0).

The phage-shaped word cloud on the landing page was generated using the ECharts wordcloud extension (v2.1.0). Genome viewers on the “Dataset Exploration” and “Genome Viewer” pages were implemented with GoslingJS (v0.11.0). On the “Data Overview” page, the Sankey diagram and pie charts were created with Apache ECharts (v5.4.0), the sunburst plot with Plotly.js (v3.0.0), and the dataset table with Tabulator (v6.3.0). All interactive plots (profile plots and heatmaps) were implemented with Plotly.js. Profile plots use fractional expression values computed from mean TPM values per gene and time point (see above). Heatmaps are based on mean TPM values and are z-score-normalized using SciPy’s `zscore` function (v1.15.2), which is also used for subsequent gene-wise clustering.

The backend was implemented in Python using Flask (v3.1.0) as a lightweight web application framework to handle API requests and URL routing. Database access is mediated via SQLAlchemy (v3.1.1), which connects the frontend to the SQLite database.

Automated accessibility checks were performed for every page according to WCAG 2.1 AA using the page scanner of the Chrome extension “Accessible Web Helper”.

The PhageExpressionAtlas runs in a Docker container hosted on the infrastructure of the Institute for Bioinformatics and Medical Informatics (IBMI) at the University of Tübingen and is integrated into the Tübingen Visualization Server (TueVis, https://tuevis.cs.uni-tuebingen.de).

### Use cases for the PhageExpressionAtlas

We conducted two use cases with the PhageExpressionAtlas web interface. For the first use case, we analyzed the dataset from [10], which profiled the dual transcriptome of T4 phage infection of *E. coli* before infection (*t* = 0) and at four time points post infection. We used the PhageExpressionAtlas to reproduce key findings from the original study, focusing on phage gene expression dynamics.

For the second use case, we analyzed Kayvirus phage infection of two *S. aureus* strains (SH1000 and Newman) from [13] by studying host ncRNA expression during phage expression, an aspect not addressed in the original publication.

### Integrative analysis with the PhageExpressionAtlas database

Beyond the use cases of the PhageExpressionAtlas web interface, we systematically queried its database allowing for integrative analysis across different phage infections.

#### Probing phage gene classification

To evaluate the classification of phage genes into early, middle and late w.r.t. infection time across diverse infections, we selected 18 datasets from 12 studies that include a pre-infection measurement and sample time points spanning the three major infection phases (Supplementary Table S1B).

For benchmarking, we defined a proxy ground truth for late gene expression based on the assumption that genes involved in virion assembly and host lysis are predominantly expressed late during infection. Accordingly, genes annotated by Pharokka as “connector”, “lysis”, “head and packaging”, “tail”, and “integration and excision” were treated as ground-truth late genes. These functions are typically enriched among late-expressed genes [8, 21]. While this proxy ground truth is necessarily imperfect and incomplete, it provides a practical benchmark for large-scale comparison across datasets. Using these labels, we assessed classification performance with the following three standard metrics.

Recall quantifies the fraction of ground-truth late genes that are predicted as late:

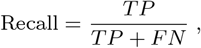

where true positives (TP) denote ground-truth late genes predicted as late, and false negatives (FN) denote ground-truth late genes not predicted as late. Under the assumption that these late labels are correct, recall represents the primary metric.

Precision measures the fraction of predicted late genes that are labeled as ground-truth late:

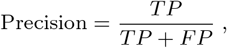

where false positives (FP) are genes predicted as late but not labeled as ground-truth late. Because the annotation-based ground truth is likely incomplete and may vary across phage taxa, we did not expect precision to be high.

As a third measure, we computed the overall fraction of genes predicted as late among all non-ground truth late genes to monitor how parameter choices affect the predicted class distribution.

We defined a grid search over (i) expression thresholds relative to each gene’s maximal expression across time points (*θ* ∈{0.1, 0.2, …, 1.0}) and (ii) relative infection-time boundaries used to bin measurements into early, middle, and late phases. For each dataset, we iterated over all parameter combinations and performed classifications using the criterion defined above. Our analysis focused primarily on the pre-computed “Class Max” and “Class Threshold” settings.

#### Expression of phage and host orthogroups and (anti-)defense systems

For the integrative analysis across infections, we selected 18 datasets from 12 studies that include a pre-infection measurement and exhibit clear transcriptional signatures of phage takeover (Supplementary Table S1B).

For all phage and host genomes, we used OrthoFinder (v3.1.2) [23] to infer orthogroups from the amino acid sequences of annotated protein-coding genes. In addition, we ran DefenseFinder (v2.0.1) [2] on the protein sequences sorted by genomic location to identify anti-phage defense systems in hosts as well as phage-encoded anti-defense systems.

For each dataset, we extracted gene expression values (fractional expression or z-score-normalized TPM means per measured time point) for genes assigned to orthogroups or to defense/anti-defense systems. Because datasets span heterogeneous time series, we harmonized infection progression by binning each time course into the following relative intervals: uninfected (pre-infection time point), immediate early (first 10%), early (subsequent 15%), middle (subsequent 25%), and late (final 50%) of the sampled infection duration. We visualized expression dynamics across datasets using boxplots (fractional expression and z-score-normalized TPM means per bin), accompanied by bar plots indicating per-dataset gene contributions. In addition, we used profile plots to display fractional expression trajectories of individual genes within each dataset.

### Results

#### Uniform processing and storage of dual RNA-seq data

We collected 42 time-resolved dual RNA-seq datasets comprising at least three infection time points from 23 peer-reviewed studies, spanning a broad range of phage-host interactions (Fig. 1A; Supplementary Table S1A). Datasets from the same study often assess the same phage infecting different host strains, the same host infected by distinct phages, or infections with phage cocktails. The largest fraction of datasets covers infections of *P. aeruginosa*, followed by *S. aureus* and *E. coli* (Fig. 1A). Overall, the collection covers transcriptomes from infections by 25 bacteriophages, predominantly virulent phages (84%) (Fig. 1B).

For some datasets, pre-computed count tables are available via GEO; however, for most studies only raw sequencing reads are accessible. We therefore established a Nextflow pipeline to process raw RNA-seq data uniformly and reproducibly into gene-level count tables under a dual-mapping setup in which reads are mapped simultaneously to the host and phage genomes (Fig. 1C). We validated the pipeline on the T4-*E. coli* dataset (comprising five time points, three biological replicates each) [10] by comparing our TPM-normalized expression matrix to the TPM-normalized matrix provided via GEO. As expected, we observed high concordance (*R*^2^ *>* 0.9) for matched samples (Supplementary Fig. S1B).

We then standardized interactive downstream processing of the count tables, which involves replicate assessment, outlier removal, and *in silico* rRNA depletion. This workflow not only provides a consistent processing strategy across studies, but also formats the data for database storage. Based on the normalized, aggregated counts tables, phage genes are assigned to the three commonly used temporal classes (“early”, “middle”, “late”) using two strategies: “Class Max” and “Class Threshold”. Replicates are aggregated for visualization within the PhageExpressionAtlas, and variance-stabilized expression values are used to select gene subsets for heatmaps. All normalized and summarized versions of the counts tables can be downloaded via the atlas.

In addition, we improved functional annotation of phage genes using Pharokka [20]. This rarely increased functional gene-level annotation, but consistently provided standardized functional categories suitable for visualization in the atlas. Finally, all processed matrices (see Methods), together with phage and host metadata and genome annotations, were stored in an SQLite database that constitutes the core data structure of the atlas backend (Fig. 1D).

We thus provide a standardized processing workflow for dual RNA-seq data of phage infections creating a unified data structure. Through application of our pipeline to 42 datasets and their storage in a database, we created the fundamental basis for the PhageExpressionAtlas.

#### Application of the PhageExpressionAtlas allows to re-assess phage-host interactions with transcriptomics

The PhageExpressionAtlas provides several functionalities for interactive interrogation of dual transcriptomes during phage infection. We use the term “atlas” by analogy to widely used single-cell transcriptomics atlases [24], which provide access to uniformly processed and integrated single-cell RNA-seq datasets and thereby serve as community “lookup” resources. Depending on the research question, users can obtain an overview of host responses and phage transcriptional dynamics (Supplementary Fig. S4, S5), classify phage genes and inspect their genomic organization (Supplementary Fig. S4, S6), or explore specific phage and host genes of interest, for example genes belonging to defined pathways or functional categories (Supplementary Fig. S5).

To illustrate these applications, we first reproduced key findings from a study characterizing the transcriptome of T4 phage infecting *E. coli* [10]. We immediately captured the dynamic transcriptional program of T4 genes over the infection time course (Fig. 2A), showing a clear separation into early, middle, and late expression classes, as reported previously [10, 22]. Inspection of phage gene classes in the genome viewer recapitulated the known clustering of temporally co-regulated genes in operon-like genomic organization as reported previously [21] (Supplementary Fig. S7A). Finally, using the “Gene Selection” panel, we revisited a canonical transcriptional cascade (Fig. 2B): three middle genes encode the proteins gp33, gp45, and gp55, which assemble into a complex that activates late transcription [25], thereby inducing late genes such as *gene 68*, which encodes the major capsid protein of bacteriophage T4 [26]. Together, this example highlights how the atlas supports exploration from global transcriptome overview and expression-based gene classification to targeted interrogation of specific regulatory modules.

**Fig. 2.**
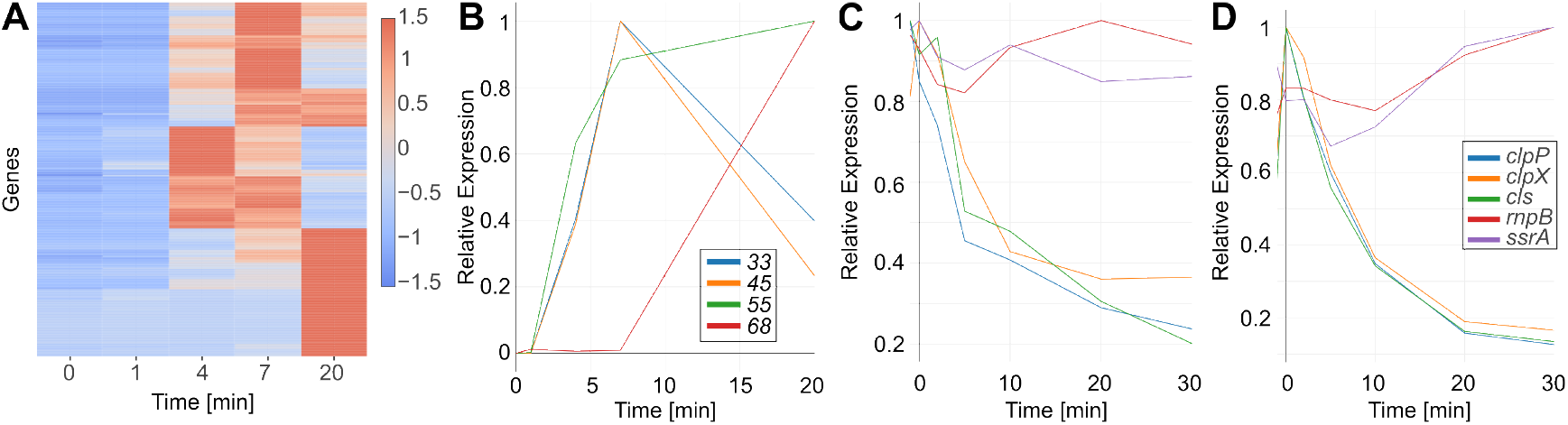
Targeted assessment of the transcriptomes of phage infections with the PhageExpressionAtlas. **A, B**) Analysis of T4 phage gene expression based on the dataset from [10]. The overall trend of phage gene expression over the time course of infection is illustrated in a z-score-normalized heatmap (**A**). Profile plots of the relative expression of the four T4 phage genes *33, 45, 55, 68*, where each gene’s expression is normalized to its maximal TPM value over the time course of infection (**B**). **C, D**) Expression of five genes (*ssrA, rnpB, clpP, clpX, cls*) in *S. aureus* strain Newman (**C**) and SH1000 (**D**) during Kayvirus phage infection [13].

The atlas provides analogous functionality for host transcriptome analysis. To demonstrate this, we revisited the observation that certain non-coding RNAs (ncRNAs), including the transfer-messenger RNA (tmRNA) SsrA and the small RNA RnpB, remain at relatively constant levels during T4 infection, suggesting that they may be host factors required for infection or otherwise maintained by the phage [10]. To assess whether this pattern generalizes, we analyzed Kayvirus infection of two *S. aureus* strains (Newman and SH1000) [13]. Using heatmaps, we recapitulated the reported global decrease in relative host transcript abundance during infection in both strains [13] and observed strain-dependent shifts in the distribution of phage gene classes (Supplementary Fig. S7B,C). While late phage genes were consistently classified as late in both hosts, infection of strain SH1000 shifted several genes classified as middle in Newman into the early class, indicating host-dependent timing differences in early and middle phage transcription.

We then examined expression profiles of the host ncRNAs *ssrA* and *rnpB* alongside essential protein-coding genes involved in proteolytic quality control (*clpP, clpX*) and lipid synthesis (*cls*). Whereas the protein-coding genes’ transcripts decreased over the infection time course in both strains, *ssrA* and *rnpB* remained comparatively abundant throughout infection (Fig. 2C,D). This observation is consistent with the hypothesis that these ncRNAs are either required during infection or are less susceptible to phage- and/or host-induced RNA turnover. Overall, these use cases illustrate how the atlas enables rapid, cross-system hypothesis testing on both the phage and host sides.

#### Assessing approaches to phage gene classification

Phage gene classification can provide a complementary route to inference of gene function, since temporal expression programs are linked to the major stages of infection progression. In the PhageExpressionAtlas, we provide two default classification schemes, “Class Max” and “Class Threshold”. “Class Max” assigns each gene to the infection phase in which it reaches its maximal expression, whereas “Class Threshold” assigns the earliest phase in which expression reaches at least 20% of the gene’s maximal value, using infection phase boundaries derived from the original studies. Such heuristic classifications are commonly used in the literature, but are rarely evaluated systematically [22, 10, 13]. We therefore leveraged the PhageExpressionAtlas database to benchmark parameter choices for expression-based phage gene classification.

We selected 18 datasets from the PhageExpressionAtlas spanning diverse phage-host interactions and covering transcriptomic measurements across the three major infection phases (early, middle, late). To quantify classification accuracy, we defined proxy ground-truth labels for a subset of late phage genes.

First, we applied this analysis to the two default schemes, “Class Max” and “Class Threshold”. “Class Max” captured a larger fraction of proxy ground-truth late genes across most datasets, whereas “Class Threshold” showed reduced recall in the majority of cases (Fig. 3A). At the same time, precision was higher for “Class Max”, indicating that a substantial fraction of genes predicted as late are represented in our annotation-derived ground-truth set. When examining the fraction of additional predicted late genes (i.e., late predictions not contained in the proxy ground truth), we observed low values for most datasets. Notably, the highest fractions occurred in phages with small (*<* 100 genes) or particularly large (*>* 300 genes) genomes (Supplementary Fig. S8A). This is consistent with sparse functional annotation in many phage genomes. We performed an additional assessment using different threshold values and time boundaries beyond our predefined classes indicating that “Class Max” represents a solid approach to phage gene classification (Supplementary Fig. S9,S10).

**Fig. 3.**
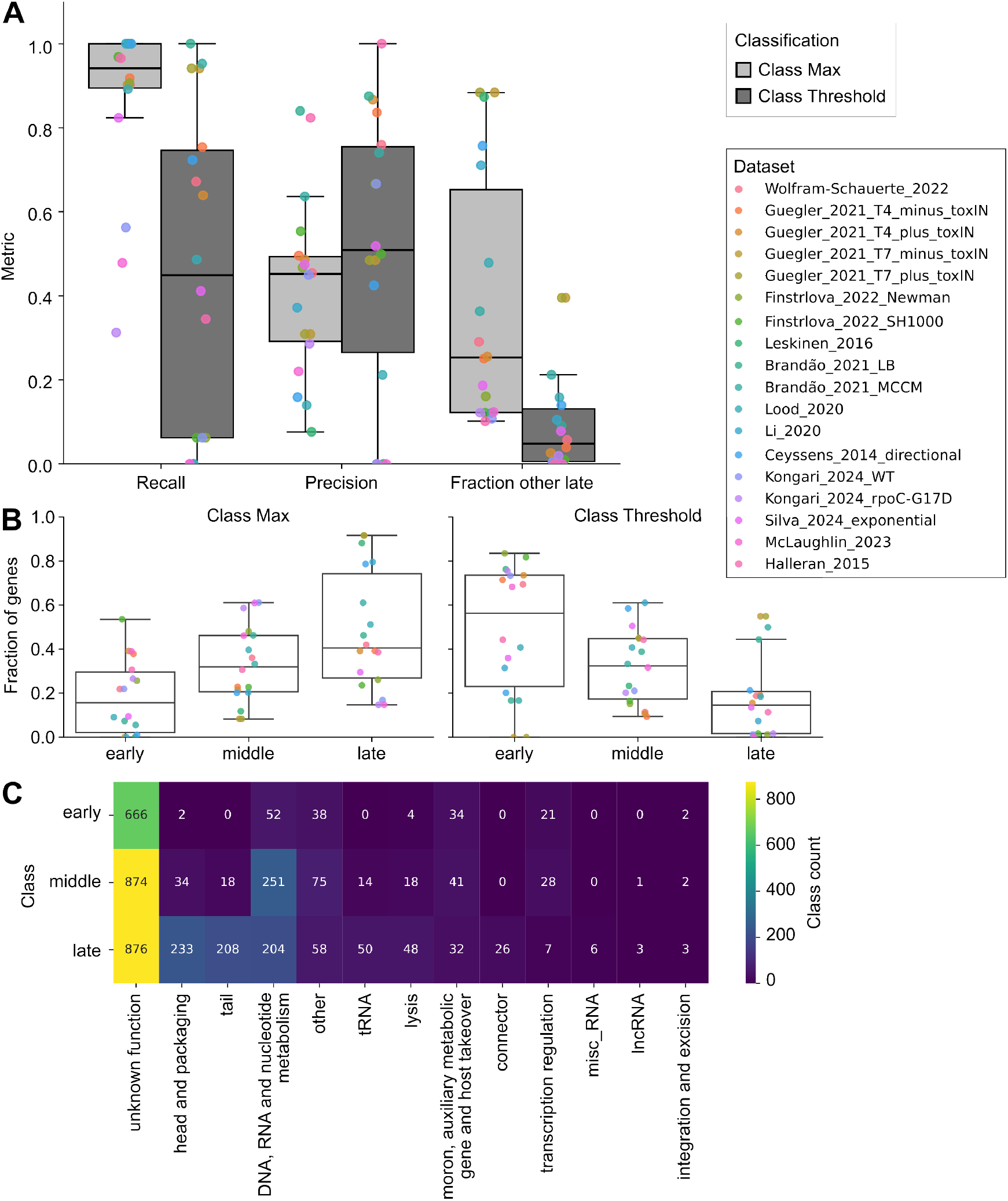
Assessment of phage gene classification methodologies. **A**) Metrics of phage gene classification for the atlas’ default settings “Class Max” and “Class Threshold” for 18 datasets. Recall defines the fraction of the Pharokka-annotated genes in the categories “connector”, “lysis”, “head and packaging”, “tail” and “integration and excision” [20] - assumed to be late genes - classified as late genes. Precision describes the fraction of predicted late genes among the presumably late genes. *Fraction* describes the percentage of predicted late genes outside the presumably late genes. **B**) Distribution of early, middle and late gene fractions after classification using “Class Max” and “Class Threshold” criteria across datasets. The dataset legend serves for A and B. **C**) Total class count using the “Class Max” criterion across Pharokka-annotated gene categories across the 18 datasets presented in A and B. Each gene is represented only once and for genes present in different datasets, majority class voting was used, retaining the earlier class in case of a tie.

We next compared the overall distribution of early, middle, and late gene fractions across datasets for both classification schemes. “Class Threshold” displayed a bias towards early classifications, whereas “Class Max” tended to yield a larger fraction of late calls (Fig. 3B; Supplementary Fig. S8B). Importantly, these distributions depend on both the sampled time points and the specific phage-host system under investigation. Notably, the same phage profiled under different growth conditions, during host manipulation (e.g., knock-down), or in distinct host strains can yield different class distributions, highlighting the sensitivity of phage transcriptional programs to environmental and host-dependent factors.

Using “Class Max”, we then assessed how predicted temporal classes distribute across functional gene groups. Despite the use of dedicated phage annotation tools [20], most phage genes remain uncharacterized and consequently dominate the early, middle, and late classes (Fig. 3C). This underscores the utility of transcriptomics-derived temporal classes as an additional layer of evidence for functional inference across the infection cycle. As expected, genes annotated as head and packaging, tail, and lysis were predominantly classified as late. Interestingly, phage-encoded tRNAs and long non-coding RNAs (lncRNAs), which are present only in a subset of phages, also tended to be classified as late, suggesting that such phage-encoded RNA-based regulatory functions may be particularly relevant at later infection stages.

Together, these analyses constitute a first cross-dataset benchmark of expression-based phage gene classification, revealing condition- and host-dependent shifts in class distributions for the same phages, as well as reproducible trends across functional gene groups.

#### The PhageExpressionAtlas reveals trends across diverse phage-host interactions

Next, we exploited the collection of time-resolved infection transcriptomes to assess recurring trends in phage and host gene expression across datasets. To this end, we selected 18 datasets that comprise at least three post-infection time points and include a transcriptome measurement of the bacterial culture prior to infection. Consistent with prior reports, the relative phage transcript abundance increases over the course of infection (Fig. 4A). Notably, the final phage transcriptome fraction varies widely between phages and datasets, ranging from below 20% to above 80% at the end of the sampled infection period, highlighting the diversity of phage-host interactions captured in the atlas.

**Fig. 4.**
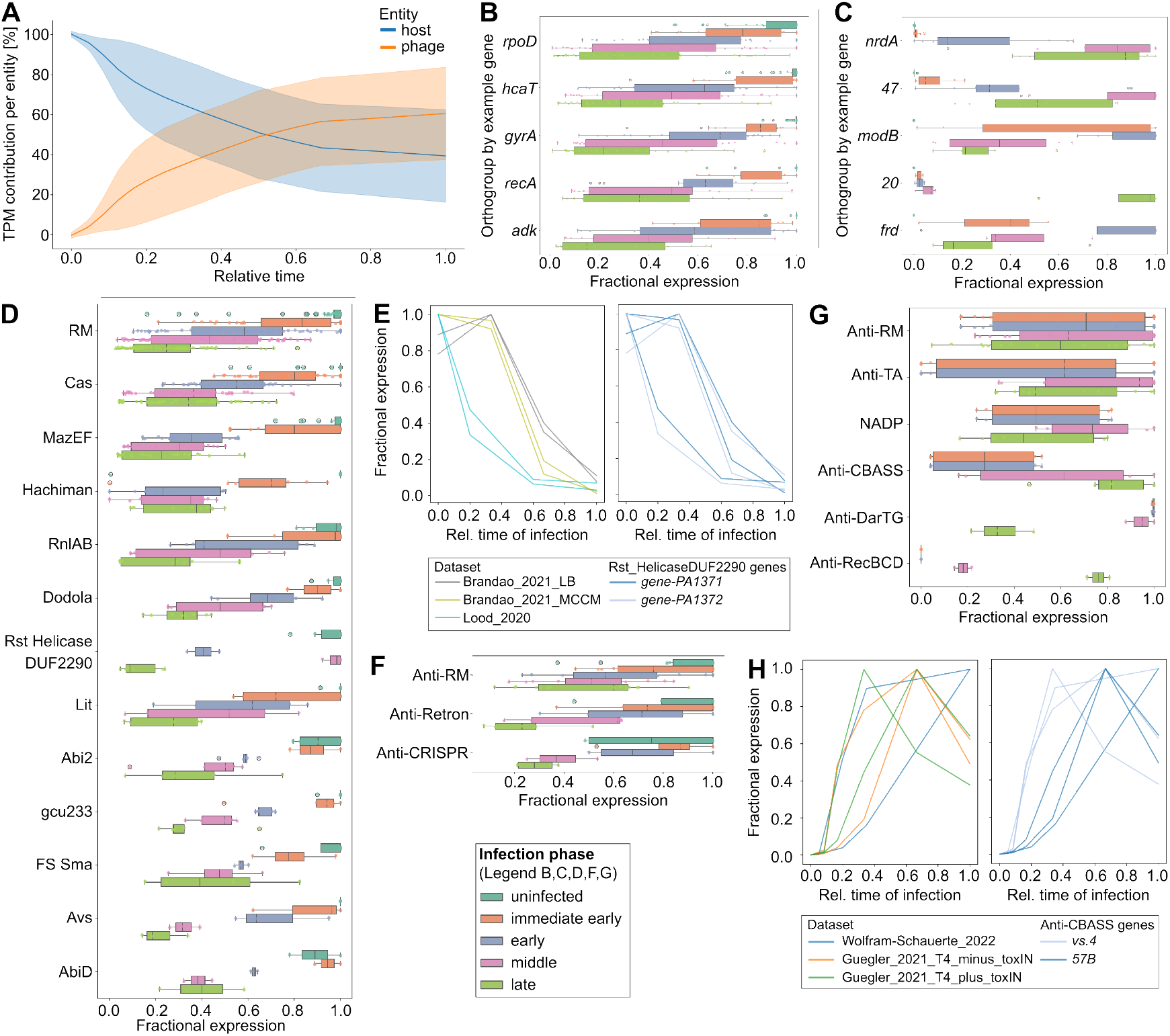
Trends of phage and host gene expression. **A**) Host and phage transcriptome abundance over the relative time course of infection. Lines represent means across datasets, the interval is determined by standard deviation from the mean (n=17 datasets). **B**) Boxplot indicating expression of orthologous genes relative to the *E. coli* house-keeping genes *rpoD, hcaT, gyrA, recA* and *adk* summarized into uninfected, immediate early, early, middle and late infection phases for n=17 datasets. Data points represent fractional expression values. Full figure in Supplementary Figure S11A. **C**) Same boxplot as in B for orthologous groups of 5 characteristic T4 phage genes. Full figure in Supplementary Figure S12A. **D**) Boxplot indicating relative expression of anti-phage defense systems in diverse bacterial hosts binned into five infection stages as in B and C. Full figure including dataset (n=17) contributions is available in Supplementary Figure S13A. **E**) Profile plot indicating relative expression of the Rst HelicaseDUF2290 defense system in *P. aeruginosa* PAO1 infected by LUZ19 (Brandao datasets) and vB PaeM PA5oct phage (Lood). **F**) Boxplot showing fractional expression of host-encoded anti-defense systems across infection phases for n=9 datasets. Full figure available in Supplementary Figure S13A. **G**) Boxplot showing fractional expression of phage-encoded anti-defense systems across immediate early, early, middle and late infection phase for n = 10 datasets. Full plot available in Supplementary Figure S14B. **H**) Profile plots illustrating relative expression of two anti-CBASS genes encoded in T4 phage across n=3 datasets. All profile plots show relative infection time on the x-axis bringing datasets to a comparable scale.

#### Expression of host and phage orthologous genes

We then used OrthoFinder [23] to infer orthogroups across bacterial hosts and across phages, respectively, to assess the expression dynamics of homologous host and phage functions across diverse infections. First, we examined orthogroups containing *E. coli* housekeeping genes, including *rpoD* (encoding the *σ*^70^ subunit of RNA polymerase), *hcaT* (a putative 3-phenylpropionate transporter), *gyrA* (DNA gyrase subunit A, essential for DNA replication), *recA* (recombinase A involved in DNA repair and the SOS response), and *adk* (adenylate kinase, central to adenylate metabolism and energy homeostasis). As expected, the relative transcript abundance of the corresponding orthologs decreases upon phage infection across a broad range of phage-host systems (Fig. 4B; Supplementary Fig. S11A). We observed a similar trend for orthogroups encoding ribosomal proteins (Supplementary Fig. S11B). Together, these patterns are consistent with a core feature of many lytic infections: progressive remodeling of transcriptome composition such that host transcripts decline in relative abundance as phage transcription increases, driven by host transcript degradation and/or dominance of the phage transcriptional program [8].

Next, we investigated orthogroups of phage genes with comparatively well-defined functions using T4 as a reference. *nrdA* encodes a ribonucleotide reductase subunit required for deoxyribonucleotide synthesis to support phage DNA replication, typically during the middle phase of infection [27, 28]. Accordingly, transcription of *nrdA* orthologs peaked predominantly in the middle phase across diverse phages infecting, among others, *P. aeruginosa, S. aureus*, and *E. coli* (Fig. 4C; Supplementary Fig. S12A). Similarly, T4 *gene 47*, which encodes a subunit of a DNA exonuclease involved in DNA repair, replication, and recombination [29], showed orthologs in multiple phages with expression maxima again concentrated in the middle phase, coinciding with the period, in which phage DNA replication typically occurs.

For canonical early genes, we found that the ADP-ribosyltransferases ModA and ModB are consistently most highly expressed in the early infection phase across datasets (Fig. 4C; Supplementary Fig. S12A). In addition, T4 *frd* transcription - with an ortholog detected in the *P. aeruginosa* phage vB PaeM PA5oct - peaked early, consistent with its role as a dihydrofolate reductase generating precursors for deoxythymidylate synthesis [30]. Conversely, T4 *gene 20*, encoding the portal protein that forms the capsid vertex required for docking of the DNA-packaging motor [31], behaved as a strictly late gene in both T4 and vB PaeM PA5oct, as expected for a structural assembly component.

Finally, screening the largest orthogroups revealed a DNA polymerase orthogroup present in six phages, with expression extending from early through middle and into early late phases depending on the phage and study (Supplementary Fig. S12B). This variability suggests that phage-encoded DNA polymerases can be deployed with different temporal programs while contributing to the shared goal of progeny production.

#### Insights into the role of defense and anti-defense systems via dual transcriptomes

Finally, we assessed the transcriptional dynamics of host-encoded defense and phage-encoded anti-defense systems across diverse phage-host interactions. This area is rapidly expanding due to the continued discovery of bacterial defense systems [2] and phage-encoded countermeasures [4]. However, their expression dynamics in time-resolved dual transcriptomes have not yet been systematically compared across “native” phage-host systems.

To this end, we employed DefenseFinder [2] to annotate host- and phage-encoded (anti-)defense systems and extracted their expression profiles from the corresponding datasets. Across multiple studies, we observed an overall decrease in relative transcript abundance for many host defense systems over the infection time course, including CRISPR-Cas, restriction-modification (RM) systems, and toxin-antitoxin (TA) modules (Fig. 4D; Supplementary Fig. S13A,B). Notably, while within-system expression profiles change over time, the rank of these systems within the host transcriptome (z-score) remains comparatively stable in many cases (Supplementary Fig. 13A,B), consistent with broad host transcriptome remodeling during lytic infection.

For a subset of systems, we observed transiently elevated expression immediately after infection. This includes the abortive infection system AbiD in a clinical *S. aureus* isolate and the Ceres defense system in *Mycobacterium smegmatis* (Fig. 4D; Supplementary Fig. S13C). These patterns suggest that some defenses may be transcriptionally induced upon infection, although transcriptional induction alone may be insufficient for effective clearance in the assayed systems. Possible explanations include limited induction magnitude, a mismatch between transcriptional and protein-level dynamics, and/or the presence of phage countermeasures.

We also observed system- and phage-dependent heterogeneity. For example, the two-gene defense system Rst HelicaseDUF2290 showed different temporal dynamics upon infection by distinct phages. During infection of *P. aeruginosa* with the large phage vB PaeM PA5oct, fractional transcript abundance decreased early, whereas during infection with the smaller phage LUZ19 (approximately 50 genes) the decrease was most apparent in the middle phase (Fig. 4E). For both phages, we did not detect annotated anti-defense systems using DefenseFinder. Given the large fraction of uncharacterized genes in vB PaeM PA5oct, additional anti-defense functions may remain unannotated, and broad RNA degradation mechanisms could also contribute to the observed dynamics, as described for T4 [32, 33].

Intriguingly, several bacterial hosts (e.g., *E. coli* and *S. aureus*) that encode multiple first-line defense systems (including RM and CRISPR-Cas) also encode corresponding anti-defense modules according to DefenseFinder predictions. In the analyzed datasets, these host-encoded anti-defense transcripts generally decreased in relative abundance over infection (Fig. 4F; Supplementary Fig. S14A), while their transcriptome rank (z-score) was often only modestly affected. This pattern suggests that substantial *de novo* transcription of these modules is not a universal hallmark of infection in the profiled conditions; functional contributions may instead depend on pre-existing protein pools, translation dynamics, or condition-specific regulation.

Finally, among the five phages in which DefenseFinder identified anti-defense systems, we examined their temporal expression programs (Fig. 4G; Supplementary Fig. S14B). Anti-RM and anti-TA modules were expressed across multiple phases, with a tendency towards earlier expression in several datasets, consistent with the expectation that nucleic-acid-targeting defenses can act rapidly after genome injection. In contrast, anti-CBASS mechanisms [34] encoded by bacteriophage T4 (e.g., *57B* /*acb1* and *vs*.*4* /*acb2*) showed predominantly middle/late expression (Fig. 4G,H), aligning with the timing of CBASS activation [35]. CBASS is triggered by cyclic nucleotides [35], which anti-CBASS proteins can degrade to prevent defense activation [34]. Notably, although CBASS is not encoded in the *E. coli* strains represented in our dataset collection, anti-CBASS expression in T4 appears as a conserved transcriptional program across conditions, potentially reflecting a broadly protective strategy rather than induction solely by the presence of CBASS.

### Discussion and Conclusion

Here, we introduce the PhageExpressionAtlas as an interactive visualization resource for exploring phage-host interactions at the transcriptional level. We established a unified processing framework to process and analyze 42 datasets from 23 studies spanning a broad range of phage-host systems. We demonstrate that the atlas enables user-friendly investigation of phage infections from global overview to in-depth analysis, including expression-based phage gene classification in genomic context. Included datasets can be re-analyzed to confirm published findings and to formulate and test new, user-defined hypotheses. To the best of our knowledge, the PhageExpressionAtlas is a first dedicated resource that provides centralized access to uniformly processed time-resolved transcriptomic data of phage infections.

Using the PhageExpressionAtlas database, we benchmarked approaches to expression-based phage gene classification. We note that this evaluation is constrained by the lack of comprehensive ground-truth labels across phages and gene classes, reflecting the large fraction of phage genes with unknown function. Nonetheless, using a proxy late-gene set enriched for structural, assembly, and lysis functions, we found that assigning temporal classes based on maximal expression (“Class Max”) correctly classifies most of these genes as late across diverse phages. Moreover, transcriptional timing was sensitive to experimental context, including growth conditions and host strain, underscoring the importance of standardized processing and cross-study comparison. Consistent with the sparse state of phage annotation, the majority of genes classified as early, middle, or late remained of unknown function, highlighting the need for continued functional characterization across infection stages and functional modules.

Future work could leverage supervised machine learning to predict phage gene classes directly from sequence, rather than from transcriptomic profiles. Such models could assist functional annotation and mechanistic investigation, as the infection phase in which a gene is expressed can be informative about the broader processes it contributes to [8]. The PhageExpressionAtlas provides a foundational training and benchmarking resource for these efforts and may complement state-of-the-art phage annotation and defense/anti-defense discovery pipelines such as Pharokka [20], Phold [36], and DefenseFinder [2].

To enable cross-system comparisons, we assessed expression of orthologous gene groups across hosts and across phages. As expected, we observed the global decline of relative host transcript abundance during lytic infection, consistent with decades of prior work [21, 10, 11, 8, 9, 37]. We recorded similar dynamics for many host-encoded defense systems. In some cases - depending on the underlying time-series resolution - defense transcripts showed an early increase immediately after infection followed by a rapid decline. Overall, these patterns suggest that defense-related transcripts are generally not selectively stabilized and may be affected by the same phage-triggered processes that remodel or degrade the host transcriptome. Previous work reported that, despite pronounced host transcriptome degradation during T4 infection, the host proteome remains comparatively stable [10]. Together, these observations are consistent with a model in which baseline (pre-infection) expression levels contribute substantially to the cell’s defense capacity, whereas *de novo* transcription during infection may play a more limited role.

In this context, a recent transcriptional survey across bacterial species and strains under diverse physiological conditions highlighted condition-dependent priming of the defense arsenal [38]. Accordingly, one plausible explanation is that many atlas datasets - often generated under near-optimal laboratory growth conditions - may capture states in which defense systems are expressed at lower levels than in more variable environmental settings, potentially lowering the barrier for successful infection even in the absence of annotated anti-defense mechanisms. At the same time, given that anti-defense discovery is still emerging, currently unannotated anti-defense modules may be identified in the datasets contained in the atlas. Moreover, a single-microbe RNA-seq-based study recently uncovered the heterogeneous expression levels of defense systems in phage-host interactions [39], which indicate that the bulk average may not accurately reflect the “native” interaction between phage and host immunity on the transcriptional level. Overall, these analyses illustrate how the PhageExpressionAtlas enables systematic, cross-dataset investigation of transcriptional dynamics of bacterial anti-phage immunity and phage counter-defense strategies, providing an entry point for mechanistic follow-up studies of this co-evolutionary arms race.

Looking forward, we plan to expand the PhageExpressionAtlas in several directions. First, we will continuously integrate additional time-resolved dual RNA-seq datasets as they become available, including datasets contributed by authors directly. Second, we will implement additional modules to further enhance interactivity and visualization, including more integrative cross-phage and cross-host analyses to enable systematic comparisons across infection systems. Finally, we aim to incorporate additional omics modalities, including proteomics [10] and transcriptome architecture data generated by differential RNA-seq [40, 41] and ONT-cappable-seq [42]. Together, these extensions would enable interrogation of multiple regulatory layers during phage infection and support a more holistic view of phage-host interactions.

## Data availability

The PhageExpressionAtlas is available at `phageexpressionatlas.cs.uni-tuebingen.de`. No novel datasets were generated in this work. All dataset accessions are documented in Supplementary Table S1A. The database is available at https://github.com/Integrative-Transcriptomics/PhageExpressionAtlas/tree/main/PhageExpressionAtlas/instance.

## Code availability

The code underlying the PhageExpressionAtlas, the creation of the database and custom analyses presented in this work is available at https://github.com/Integrative-Transcriptomics/PhageExpressionAtlas and archived in zenodo at https://doi.org/10.5281/zenodo.19249623. The processing pipeline is available at https://github.com/Integrative-Transcriptomics/phage_host_dual_transcriptomics and archived in zenodo at https://doi.org/10.5281/zenodo.19249672.

## Supplementary Data

Supplementary figures are available in the Supplementary Information. Supplementary Tables S1A and S1B are available via Supplementary Table S1.xlsx.

## Author contributions statement

M.W.-S.: writing and conceptualization, funding acquisition, frontend and backend development, data analysis; N.W.: backend development and data analysis; C.T.: frontend development; K.N.: writing and conceptualization, funding acquisition.

## Acknowledgments

This work was supported by funding from the Wilhelm Schuler Stiftung (University of Tübingen) awarded to M.W.-S. for the development of the PhageExpressionAtlas. We acknowledge the user feedback on the PhageExpressionAtlas from former and current colleagues.

## References

1. L. O’Sullivan, C. Buttimer, O. McAuliffe, D. Bolton, and Coffey. Bacteriophage-based tools: recent advances and novel applications. F1000Res, 5:2782, 2016.

2. F. Tesson, A. Herve, E. Mordret, M. Touchon, C. d’Humieres, J. Cury, and A. Bernheim. Systematic and quantitative view of the antiviral arsenal of prokaryotes. Nat Commun, 13(1):2561, 2022.

3. J. Y. Wang and J. A. Doudna. Crispr technology: A decade of genome editing is only the beginning. Science, 379(6629):eadd8643, 2023.

4. K. Murtazalieva, A. Mu, A. Petrovskaya, and R. D. Finn. The growing repertoire of phage anti-defence systems. Trends Microbiol, 32(12):1212–1228, 2024.

5. M. Gerovac, K. Chihara, L. Wicke, B. Bottcher, R. Lavigne, and J. Vogel. Phage proteins target and co-opt host ribosomes immediately upon infection. Nat Microbiol, 9(3):787–800, 2024.

6. M. Wolfram-Schauerte, N. Pozhydaieva, J. Grawenhoff, L. M. Welp, I. Silbern, A. Wulf, F. A. Billau, T. Glatter, H. Urlaub Jaschke, and K. Hofer. A viral adp-ribosyltransferase attaches rna chains to host proteins. Nature, 620(7976):1054–1062, 2023.

7. M. Skurnik, S. Alkalay-Oren, M. Boon, M. Clokie, T. Sicheritz-Pontén, K. Dabrowska, G. F. Hatfull, R. Hazan, M. Jalasvuori, S. Kiljunen, R. Lavigne, D. J. Malik, R. Nir-Paz, and J.-P. Pirnay. Phage therapy. Nature Reviews Methods Primers, 5(1):9, 2025.

8. L. Putzeys, L. Wicke, A. Brandao, M. Boon, D. P. Pires, J. Azeredo, J. Vogel, R. Lavigne, and M. Gerovac. Exploring the transcriptional landscape of phage-host interactions using novel high-throughput approaches. Curr Opin Microbiol, 77:102419, 2024.

9. A. Brandao, D. P. Pires, L. Coppens, M. Voet, R. Lavigne, and J. Azeredo. Differential transcription profiling of the phage luz19 infection process in different growth media. RNA Biol, 18(11):1778–1790, 2021.

10. M. Wolfram-Schauerte, N. Pozhydaieva, M. Viering, T. Glatter, and K. Höfer. Integrated omics reveal time-resolved insights into t4 phage infection of e. coli on proteome and transcriptome levels. Viruses, 14(11), 2022.

11. C. K. Guegler and M. T. Laub. Shutoff of host transcription triggers a toxin-antitoxin system to cleave phage rna and abort infection. Mol Cell, 81(11):2361–2373 e9, 2021.

12. C. Lood, K. Danis-Wlodarczyk, B. G. Blasdel, H. B. Jang, D. Vandenheuvel, Y. Briers, J. P. Noben, V. van Noort, Z. Drulis-Kawa, and R. Lavigne. Integrative omics analysis of pseudomonas aeruginosa virus pa5oct highlights the molecular complexity of jumbo phages. Environ Microbiol, 22(6):2165–2181, 2020.

13. A. Finstrlova, I. Maslanova, B. G. Blasdel Reuter, J. Doskar, F. Gotz, and R. Pantucek. Global transcriptomic analysis of bacteriophage-host interactions between a kayvirus therapeutic phage and staphylococcus aureus. Microbiol Spectr, 10(3):e0012322, 2022.

14. R. H. Wang, S. Yang, Z. Liu, Y. Zhang, X. Wang, Z. Xu, J. Wang, and S. C. Li. Phagescope: a well-annotated bacteriophage database with automatic analyses and visualizations. Nucleic Acids Res, 52(D1):D756–D761, 2024.

15. D. A. Russell and G. F. Hatfull. Phagesdb: the actinobacteriophage database. Bioinformatics, 33(5):784–786, 2017.

16. P. Di Tommaso, M. Chatzou, E. W. Floden, P. P. Barja, E. Palumbo, and C. Notredame. Nextflow enables reproducible computational workflows. Nat Biotechnol, 35(4):316–319, 2017.

17. S. Andrews. Fastqc: A quality control tool for high throughput sequence data, 2010.

18. M. Martin. Cutadapt removes adapter sequences from high-throughput sequencing reads. EMBnet.journal, 17(1):10–12, 2011.

19. D. Kim, J. M. Paggi, C. Park, C. Bennett, and S. L. Salzberg. Graph-based genome alignment and genotyping with hisat2 and hisat-genotype. Nat Biotechnol, 37(8):907–915, 2019.

20. G. Bouras, R. Nepal, G. Houtak, A. J. Psaltis, P. J. Wormald, and S. Vreugde. Pharokka: a fast scalable bacteriophage annotation tool. Bioinformatics, 39(1), 2023.

21. E. S. Miller, E. Kutter, G. Mosig, F. Arisaka, T. Kunisawa, and W. Rüger. Bacteriophage t4 genome. Microbiol Mol Biol Rev, 67(1):86–156, table of contents, 2003.

22. K. Luke, A. Radek, X. Liu, J. Campbell, M. Uzan, R. Haselkorn, and Y. Kogan. Microarray analysis of gene expression during bacteriophage t4 infection. Virology, 299(2):182–91, 2002.

23. D. M. Emms and S. Kelly. Orthofinder: phylogenetic orthology inference for comparative genomics. Genome Biol, 20(1):238, 2019.

24. A. Regev, S. A. Teichmann, E. S. Lander, I. Amit, C. Benoist, E. Birney, B. Bodenmiller, P. Campbell, P. Carninci, M. Clatworthy, H. Clevers, B. Deplancke, I. Dunham, J. Eberwine, R. Eils, W. Enard, A. Farmer, L. Fugger, B. Gottgens, N. Hacohen, M. Haniffa, M. Hemberg, S. Kim, P. Klenerman, A. Kriegstein, E. Lein, S. Linnarsson, E. Lundberg, J. Lundeberg, P. Majumder, J. C. Marioni, M. Merad, M. Mhlanga, M. Nawijn, M. Netea, G. Nolan, D. Pe’er, A. Phillipakis, C. P. Ponting, S. Quake, W. Reik, O. Rozenblatt-Rosen, J. Sanes, R. Satija, T. N. Schumacher, A. Shalek, E. Shapiro, P. Sharma, J. W. Shin, O. Stegle, M. Stratton, M. J. T. Stubbington, F. J. Theis, M. Uhlen, van Oudenaarden, A. Wagner, F. Watt, J. Weissman Wold, R. Xavier, N. Yosef, and Participants Human Cell Atlas Meeting. The human cell atlas. Elife, 6, 2017.

25. S. Nechaev, M. Kamali-Moghaddam, E. Andre, J. P. Leonetti, and E. P. Geiduschek. The bacteriophage t4 late-transcription coactivator gp33 binds the flap domain of escherichia coli rna polymerase. Proc Natl Acad Sci U S A, 101(50):17365–70, 2004.

26. B. Keller, C. Sengstag, E. Kellenberger, and T. A. Bickle. Gene 68, a new bacteriophage t4 gene which codes for the 17k prohead core protein is involved in head size determination. J Mol Biol, 179(3):415–30, 1984.

27. M. J. Tseng, P. He, J. M. Hilfinger, and G. R. Greenberg. Bacteriophage t4 nrda and nrdb genes, encoding ribonucleotide reductase, are expressed both separately and coordinately: characterization of the nrdb promoter. J Bacteriol, 172(11):6323–32, 1990.

28. M. J. Tseng, J. M. Hilfinger, P. He, and G. R. Greenberg. Tandem cloning of bacteriophage t4 nrda and nrdb genes and overproduction of ribonucleoside diphosphate reductase (alpha 2 beta 2) and a mutationally altered form (alpha 2 beta 2(93)). J Bacteriol, 174(17):5740–4, 1992.

29. T. J. Herdendorf, D. W. Albrecht, S. J. Benkovic, and S. W. Nelson. Biochemical characterization of bacteriophage t4 mre11-rad50 complex. J Biol Chem, 286(4):2382–92, 2011.

30. R. A. Mosher, A. B. DiRenzo, and C. K. Mathews. Bacteriophage t4 virion dihydrofolate reductase: approaches to quantitation and assessment of function. J Virol, 23(3):645–58, 1977.

31. L. Sun, X. Zhang, S. Gao, P. A. Rao, V. Padilla-Sanchez, Z. Chen, S. Sun, Y. Xiang, S. Subramaniam, V. B. Rao, and M. G. Rossmann. Cryo-em structure of the bacteriophage t4 portal protein assembly at near-atomic resolution. Nat Commun, 6:7548, 2015.

32. M. Uzan. Rna processing and decay in bacteriophage t4. Prog Mol Biol Transl Sci, 85:43–89, 2009.

33. M. Uzan and E. S. Miller. Post-transcriptional control by bacteriophage t4: mrna decay and inhibition of translation initiation. Virol J, 7:360, 2010.

34. S. J. Hobbs, T. Wein, A. Lu, B. R. Morehouse, J. Schnabel Leavitt, E. Yirmiya, R. Sorek, and P. J. Kranzusch. Phage anti-cbass and anti-pycsar nucleases subvert bacterial immunity. Nature, 605(7910):522–526, 2022.

35. D. Cohen, S. Melamed, A. Millman, G. Shulman, Y. Oppenheimer-Shaanan, A. Kacen, S. Doron, G. Amitai, and R. Sorek. Cyclic gmp-amp signalling protects bacteria against viral infection. Nature, 574(7780):691–695, 2019.

36. G. Bouras, S. R. Grigson, M. Mirdita, M. Heinzinger, B. Papudeshi, V. Mallawaarachchi, R. Green, R. S. Kim, V. Mihalia, A. J. Psaltis, P. J. Wormald, S. Vreugde, M. Steinegger, and R. A. Edwards. Protein structure-informed bacteriophage genome annotation with phold. Nucleic Acids Res, 54(1), 2026.

37. Z. Yang, S. Yin, G. Li, J. Wang, G. Huang, B. Jiang, B. You, Y. Gong, C. Zhang, X. Luo, Y. Peng, and X. Zhao. Global transcriptomic analysis of the interactions between phage phiabp1 and extensively drug-resistant acinetobacter baumannii. mSystems, 4(2), 2019.

38. L. Paoli, B. Laruelle, R. Lavenir, A. Loubat, F. Tesson, B. Gaborieau, and A. Bernheim. Environment and physiology shape antiphage system expression. bioRxiv, 2025.

39. A. Gupta, N. Morella, D. Sutormin, N. Li, K. Gaisser, Robertson, Y. Ispolatov, G. Seelig, N. Dey, and A. Kuchina. Dynamics of phage-host interactions in bacteroides fragilis resolved by single-cell transcriptomics. Nat Commun, 2026.

40. L. Wicke, F. Ponath, L. Coppens, M. Gerovac, R. Lavigne, and J. Vogel. Introducing differential rna-seq mapping to track the early infection phase for pseudomonas phage ϕkz. RNA Biol, 18(8):1099–1110, 2021.

41. M. Wolfram-Schauerte, A. Moskalchuk, N. Pozhydaieva, A. R. Rojas, D. Schindler, S. Kaiser, N. Pazcia, and K. Höfer. T4 phage rna is nad-capped and alters the nad-cap epitranscriptome of escherichia coli during infection through a phage-encoded decapping enzyme. bioRxiv, page 2024.04.04.588121, 2024.

42. L. Putzeys, M. Boon, E. M. Lammens, K. Kuznedelov, K. Severinov, and R. Lavigne. Development of ont-cappable-seq to unravel the transcriptional landscape of pseudomonas phages. Comput Struct Biotechnol J, 20:2624–2638, 2022.

