## Supplementary Information for "The PhageExpressionAtlas reveals shared and unique transcriptional patterns across phage-host interactions"

March 2026

### 1 Supplementary Figures

- Supplementary Fig. S1 presents complimentary information on datasets available in the PhageExpressionAtlas and Nextflow pipeline validation.
- Supplementary Fig. S2 - S6 provide additional illustrations of the interface of the PhageExpressionAtlas.
- Supplementary Fig. S7 shows use cases of the Genome Viewer for phage gene classification in the genomic context.
- Supplementary Fig. S8, S9 and S10 show additional information regarding the benchmarking and analysis of phage gene classification.
- Supplementary Fig. S11 - S14 present additional data for cross-host and phage expression analysis including defense and anti-defense systems.

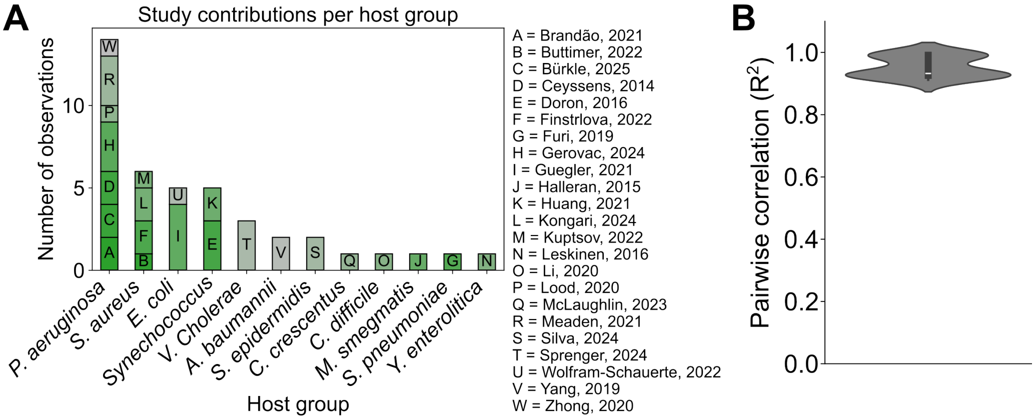


Supplementary Figure S1: **Data overview and pipeline validation.**
**A**) Barplot representing the number of datasets per bacterial host available in the PhageExpressionAtlas and the contribution of datasets by individual studies (n=42). **B**) Violinplot illustrating the pairwise $R^2$ value as a quality metric for the bulk RNA-seq data processed with the Nextflow pipeline used for data available in the PhageExpressionAtlas compared to the data provided on GEO. The comparison has been performed after in silico rRNA depletion and TPM normalization including all remaining phage and host genes for the Wolfram-Schauerte_2022 dataset [1]. Sample-wise comparisons clearly indicate that the Nextflow pipeline is able to reproduce previously processed bulk RNA-seq data.

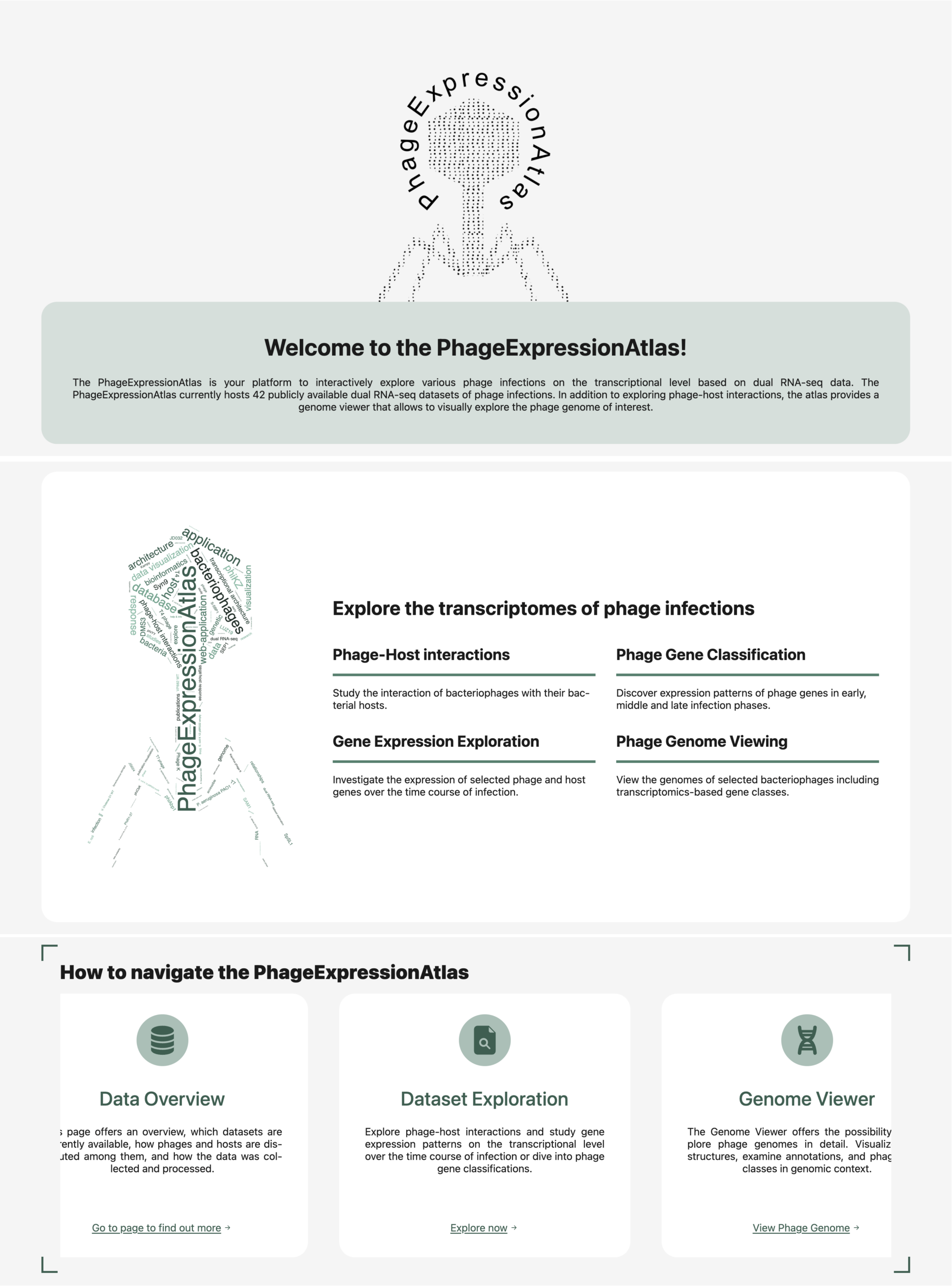


Supplementary Figure S2: **Landing page of the PhageExpressionAtlas.**
Excerpts from the PhageExpressionAtlas landing page representing the main elements on the initial landing page providing information on the atlas and the related topic (dual transcriptomics of phage infections).

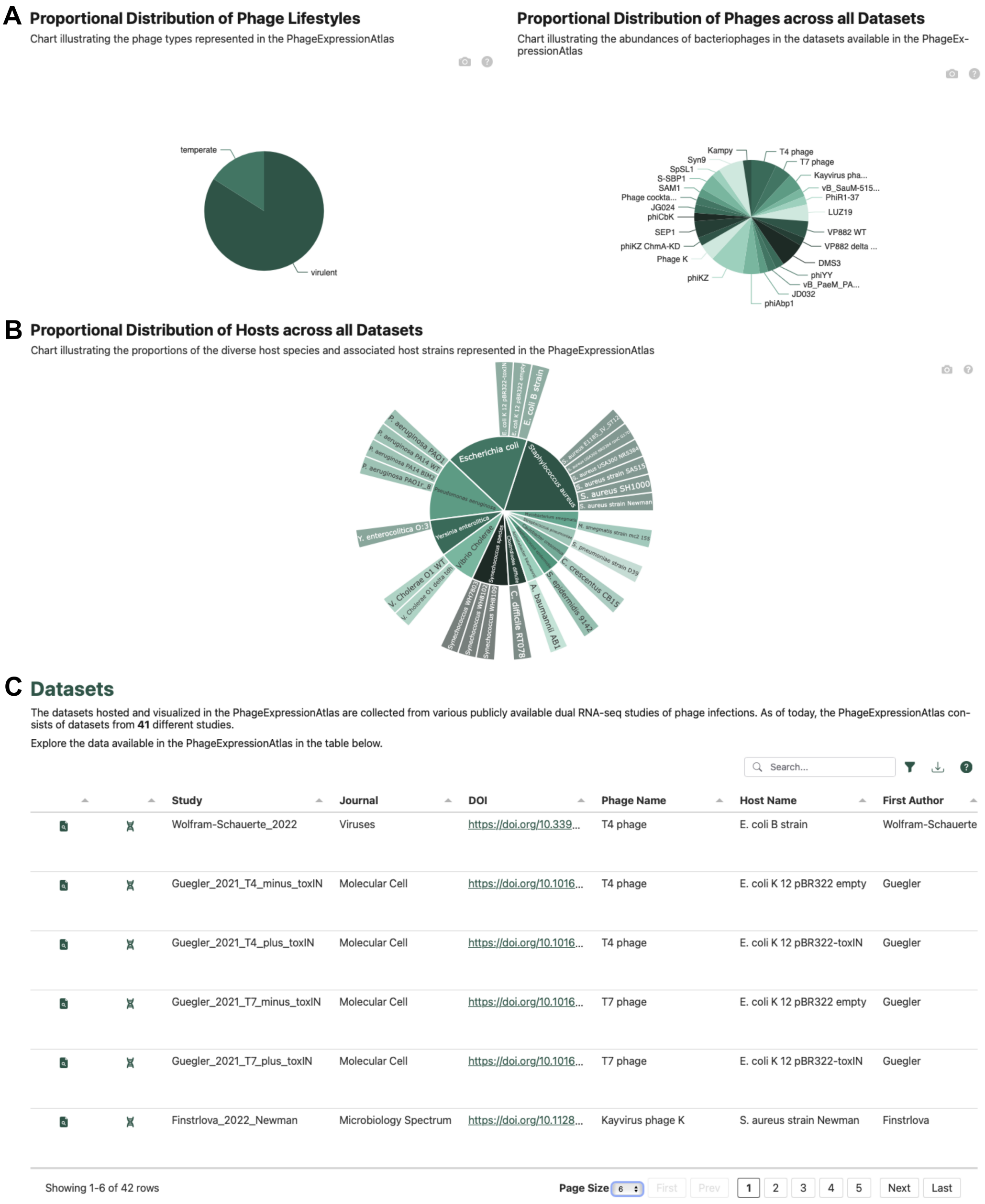


Supplementary Figure S3: **Features from the Data Overview page of the PhageExpressionAtlas.** The Data Overview page provides visualizations of data distribution in the database of the PhageExpressionAtlas including phage lifestyles and proportional distribution of phages across the datasets (**A**) as well as proportional distribution of host species and strains in a Sunburst plot (**B**). Further, a table allows to interactively search for datasets of interest linking to the Dataset Exploration and Genome Viewer subpages (**C**).

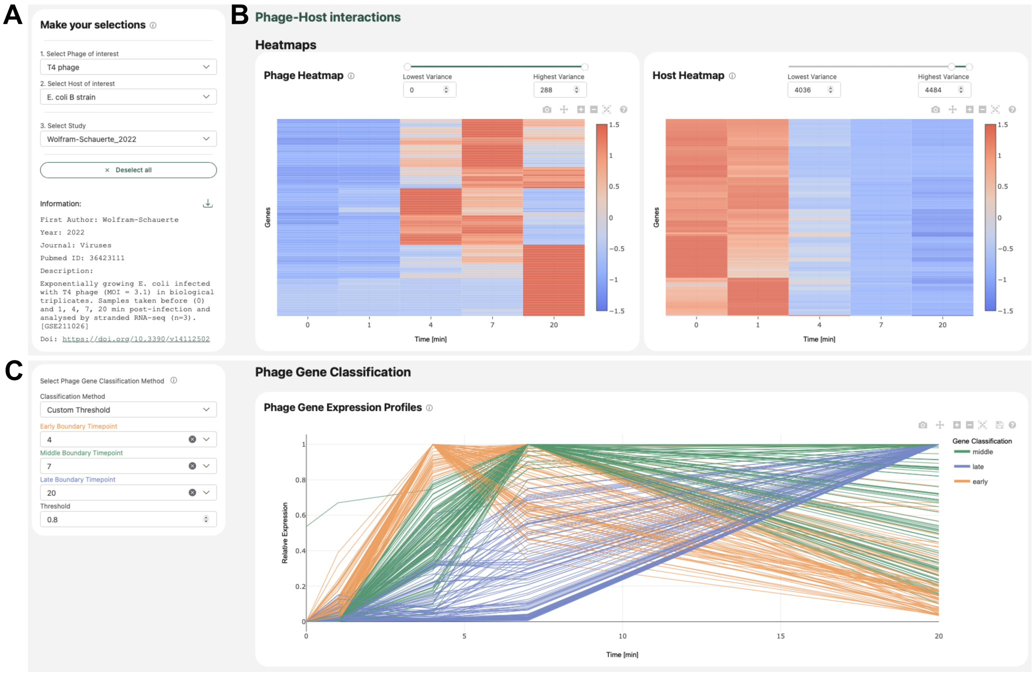


Supplementary Figure S4: **Interactive visual exploration of phage and host gene expression with the PhageExpressionAtlas.**
Representative excerpts from the "Dataset Exploration' page from the PhageExpressionAtlas. **A**) First, a dataset is selected with the dropdown menu on the left-hand side including phage, host and dataset. Corresponding dataset details are provided below. **B**) On the right-hand side, two heatmap visualizations of phage and host gene expression provide an overview of overall expression patterns. For this, replicates are summarized after TPM-normalization and z-score-normalized. All phage genes are displayed and the same number of most variable host genes is initially shown in the right-hand heatmap. The number of genes to be plotted can be selected with the variance filters. **C**) Below, phage genes can be classified into the classes "early', "middle' and "late'. For this, one of three options - either the pre-defined "Class Max' or "Class Threshold' or the "Custom Threshold' (shown) can be selected and the classification is instantly shown on the right-hand-side profile plot including all expressed phage genes. Fractional expression of replicates summarized per time point are the basis for the visualization. All examples display expression data for T4 phage infection of *E. coli* B strain [1].

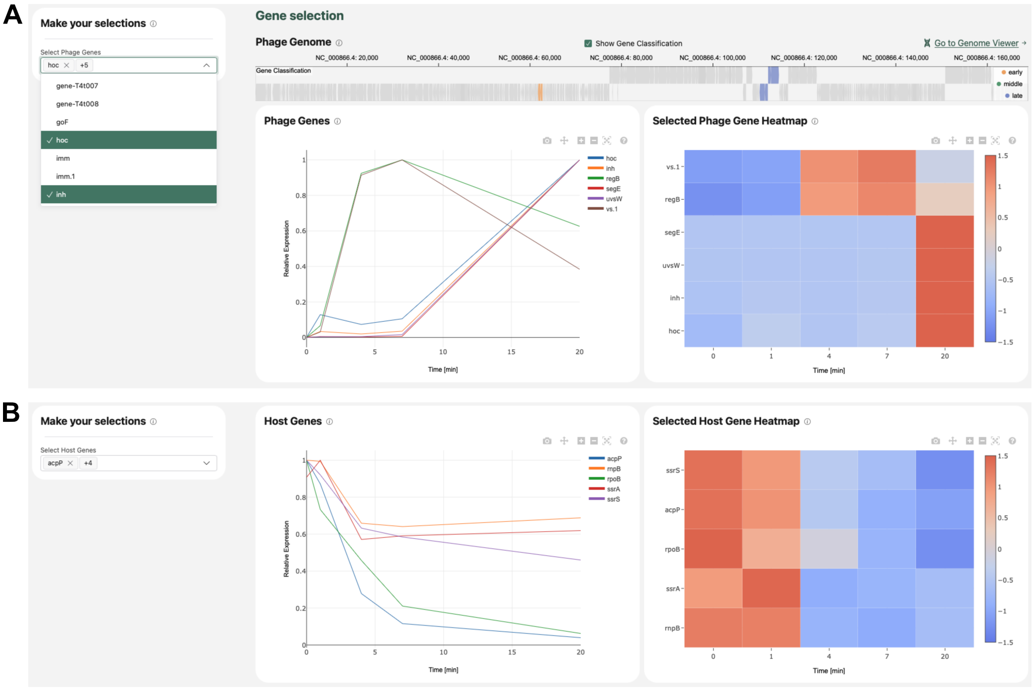


Supplementary Figure S5: **Phage and host gene expression analysis with the PhageExpressionAtlas. A**) Visualization of expression of a subset of T4 phage genes, given a selection of genes from a searchable dropdown menu, in a profile plot and heatmap and their highlighting in phage genomic context above. **B**) Exemplary investigation of the expression of the genes *ssrA*, *ssrS*, *rnpB*, *rpoB* and *acpP* in *E. coli* infected by T4 phage in the dataset from [1]. The data represents an excerpt from the PhageExpressionAtlas subpage Dataset Exploration.

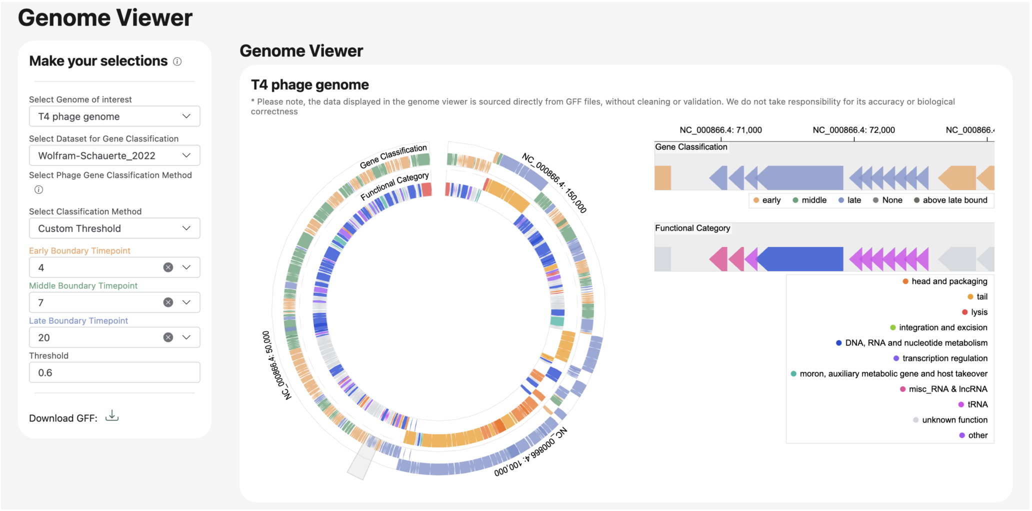


Supplementary Figure S6: **Exploration of phage gene function and class in the genomic context.** Users can select the phage genome of their interest, a corresponding dual RNA-seq dataset and the classification method. A dual-layer genome ring displays the entire phage genome in a circular manner with the gene's functional category (e.g. DNA, RNA and nucleotide metabolism) in the inner ring and the classification in the outer ring. Specific regions of the phage genome can be zoomed into with the selection window and a linear view of this selection is provided on the right-hand side. The provided example displays the T4 phage genome classified based on the dataset from [1] using the Custom Threshold method.

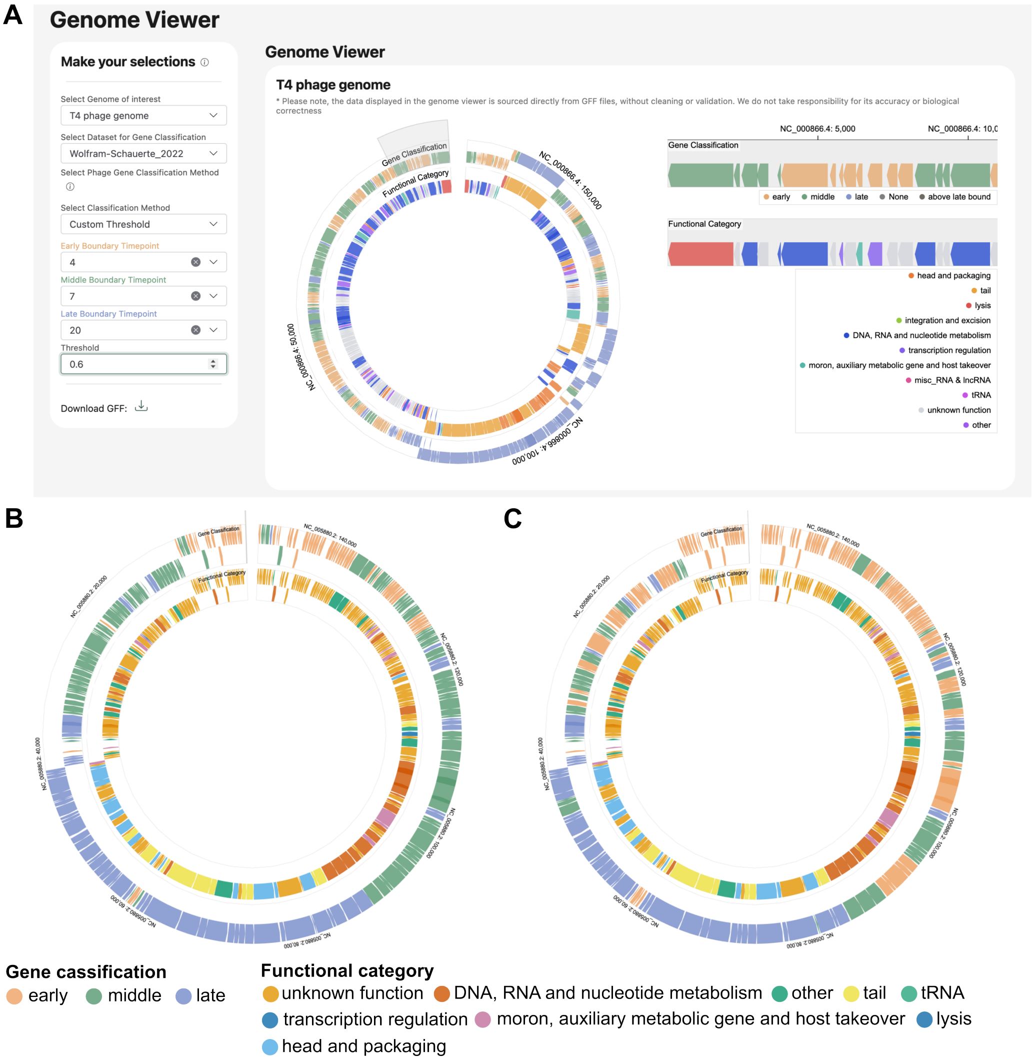


Supplementary Figure S7: **Use cases of the Genome Viewer of the PhageExpressionAtlas.** **A**) T4 phage gene classification with the custom threshold (0.6) in the Genome Viewer on the basis of the Wolfram-Schauerte_2022 dataset [1]. Phage gene classes form clear genomic clusters with late genes involved in lysis and phage assembly predominantly occupying the center of the genome. **B, C**) Comparative gene classification in phage K based on infection of S. aureus strain Newman (B) or SH1000 (C) based on data from Finstrlova-2022 [2]. The color legend for gene class (outer ring) and functional category (inner ring) is valid for B and C. Classification was performed with a custom theshold of 1 using 10 min as early, 20 min as middle and 30 min as late boundary. Clearly, the host strain seems to affect predominantly early and middle phage gene classes. All representations are screenshots from the PhageExpressionAtlas.

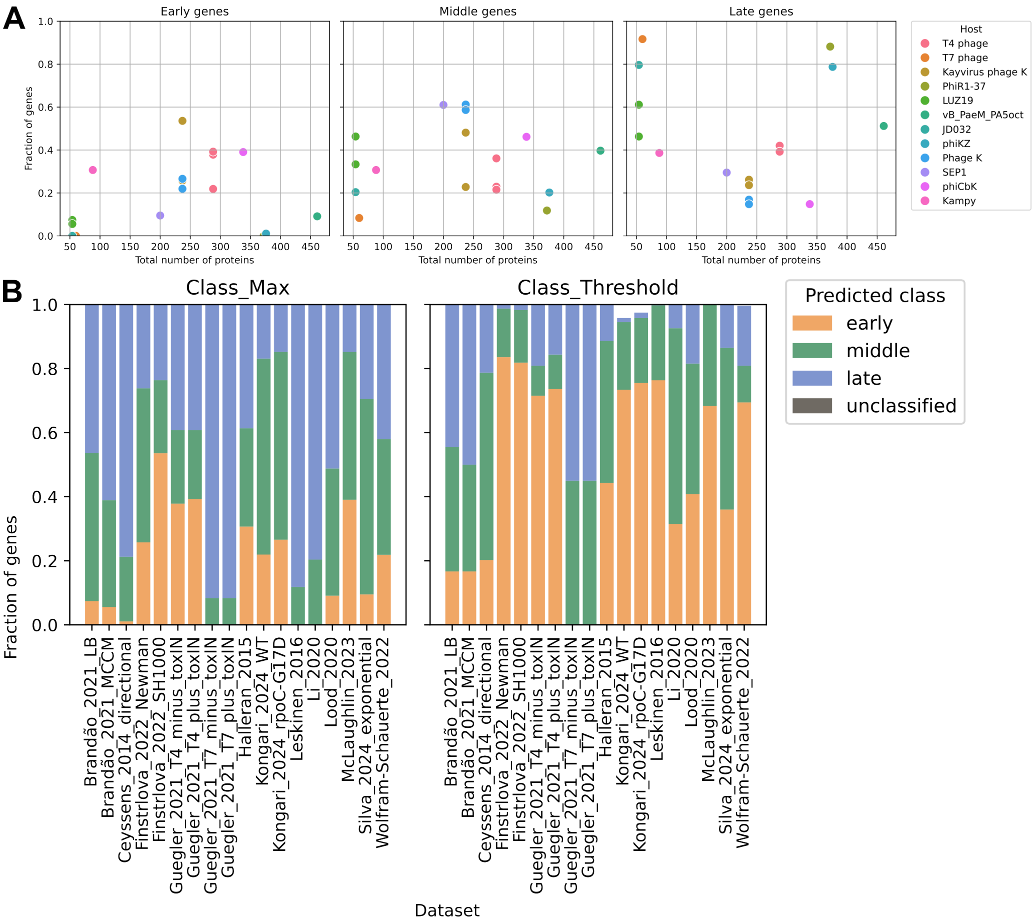


Supplementary Figure S8: **Phage gene classifications with respect to phage genome and datasets.** **A**) Phage gene classification across 18 datasets with Class Max plotted against the number of genes in the respective phage. Some phages are represented several times, since they are available in different datasets. Class Max seems to assign comparably high fractions of late genes to phages with very few and comparably many genes in total. **B**) Distribution of early, middle and late genes across 18 datasets using Class Max (left) and Class Threshold (right).

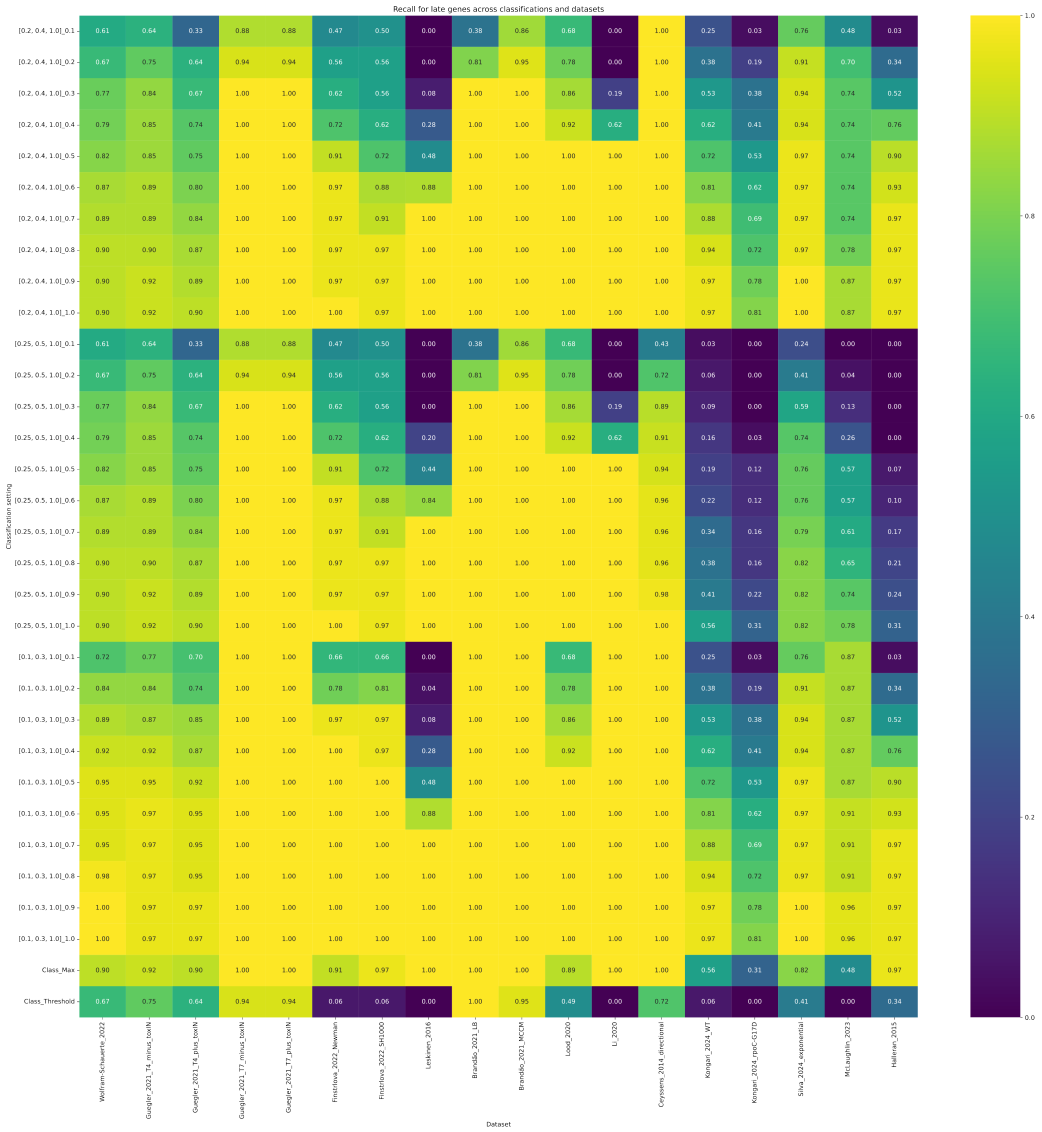


Supplementary Figure S9: **Screening of phage gene classification parameters via recall.** Recall of late phage gene classification across various classification parameters. Recall is calculated as the fraction of ground-truth late genes (defined by functions such as connector, tail, lysis, head and packaging) correctly classified as late. Recall has been assessed for 18 datasets across the default settings Class Max and Class Threshold as well as mutliple other settings. Here, relative time boundaries are given as [early, middle, late] together with a _threshold for classification.

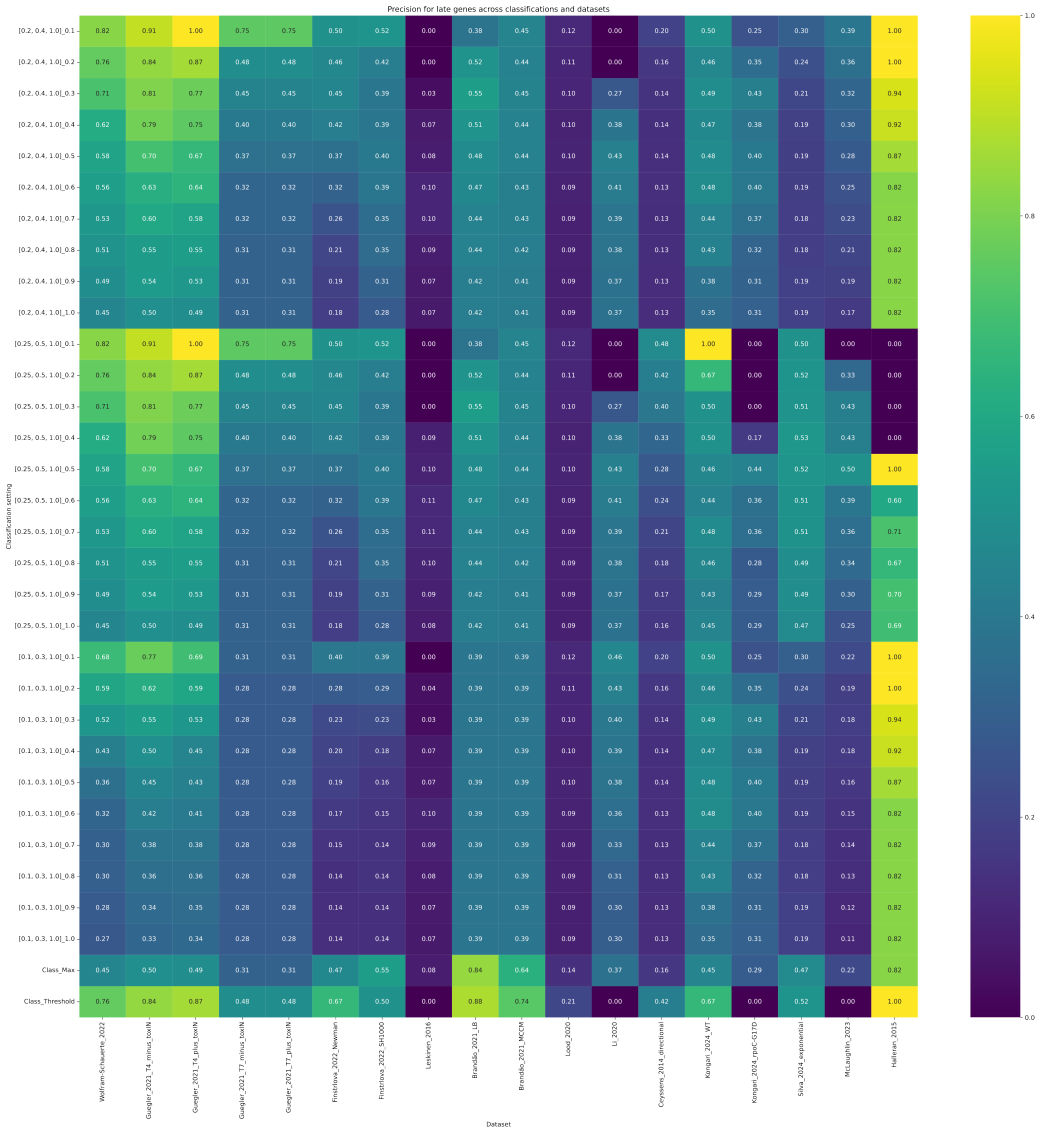


Supplementary Figure S10: **Screening of phage gene classification parameters via precision.** Precision of late phage gene classification across various classification parameters. Precision is calculated as the fraction of classified late genes that are labelled as ground-truth late genes (defined by functions such as connector, tail, lysis, head and packaging). Precision has been assessed for 18 datasets across the default settings Class Max and Class Threshold as well as mutliple other settings. Here, relative time boundaries are given as [early, middle, late] together with a _threshold for classification.

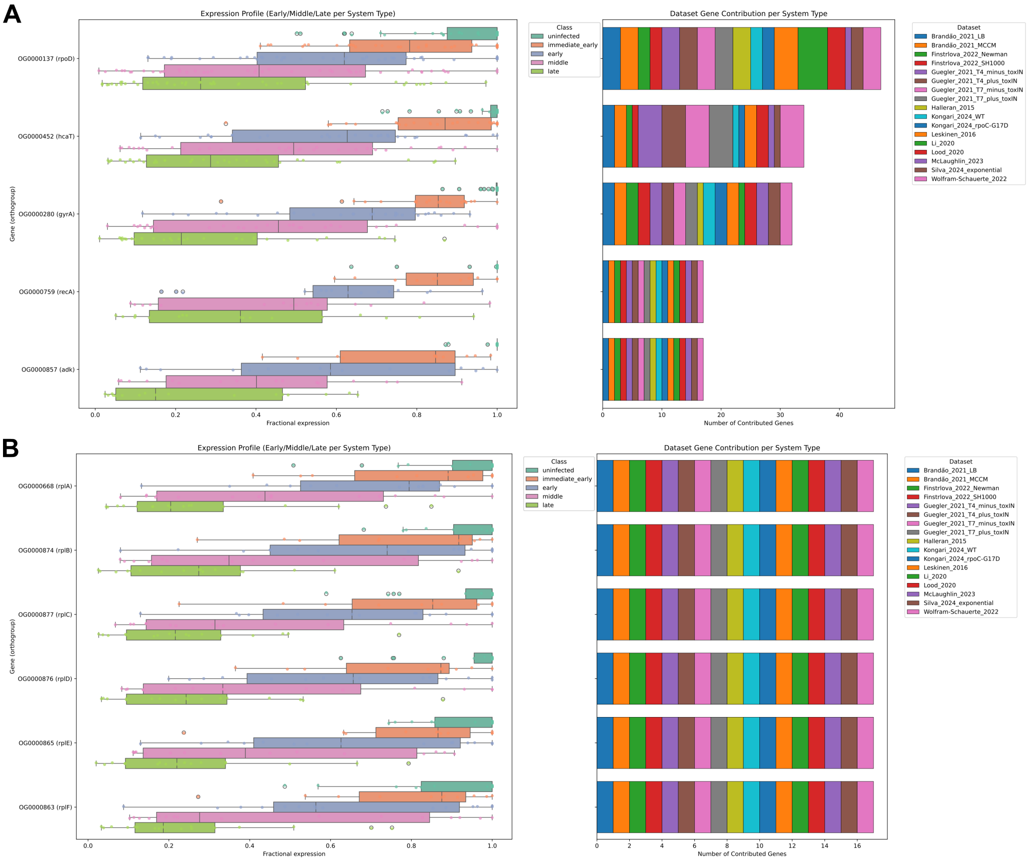


Supplementary Figure S11: **Characterization of expression of orthologous host genes during phage infection.**

**A**) Fractional expression of orthologous groups including *E. coli* house-keeping genes (*rpoD*, *hcaT*, *gyrA*, *recA*, *adk*) in uninfected hosts as well as in immediate early, early, middle and late infection phase (left). Stacked bar plot (right) indicates the number of genes contributed to each orthogroup by the individual datasets. **B**) Same plot as for the house-keeping genes for ribosomal protein-coding genes *rplA* - *rplF*. All data is sampled from 17 datasets in total.

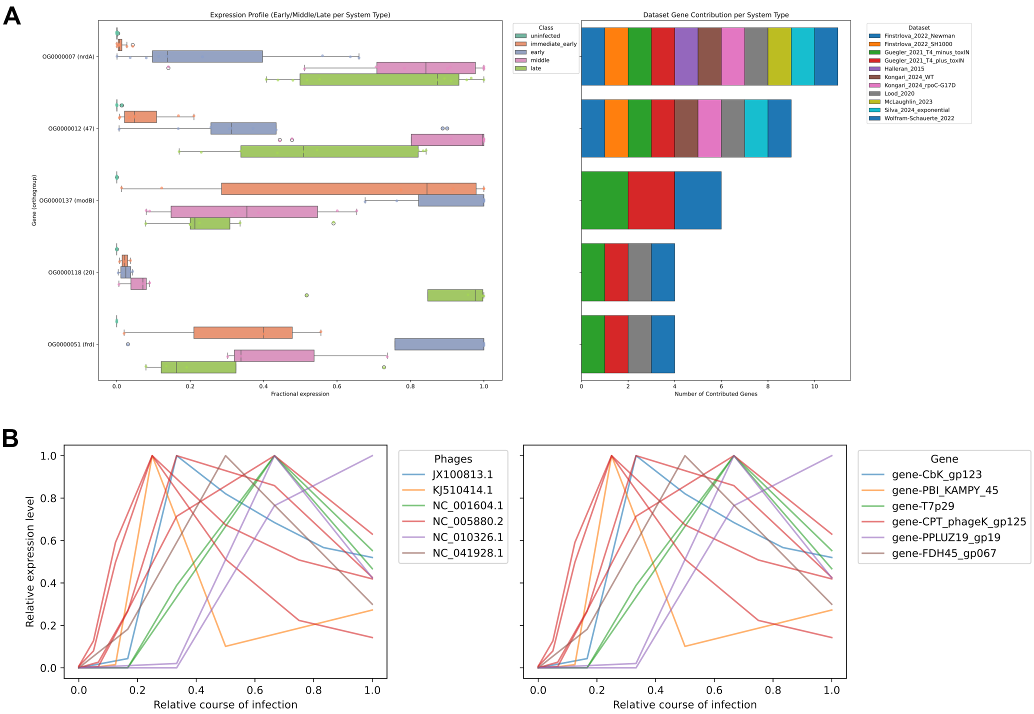


Supplementary Figure S12: **Characterization of expression of orthologous phage genes during infection.**

**A**) Fractional expression of orthologous groups including characteristic T4 phage genes (*nrdA*, *47*, *modB*, *20*, *frd*) in uninfected hosts as well as in immediate early, early, middle and late infection phase (left). Stacked bar plot (right) indicates the number of genes contributed to each orthogroup by the individual datasets. **B**) Profile plot indicating the expression of phage DNA polymerases included in a group of orthologous genes colored by associated phage and gene. All data is subsampled from 17 datasets.

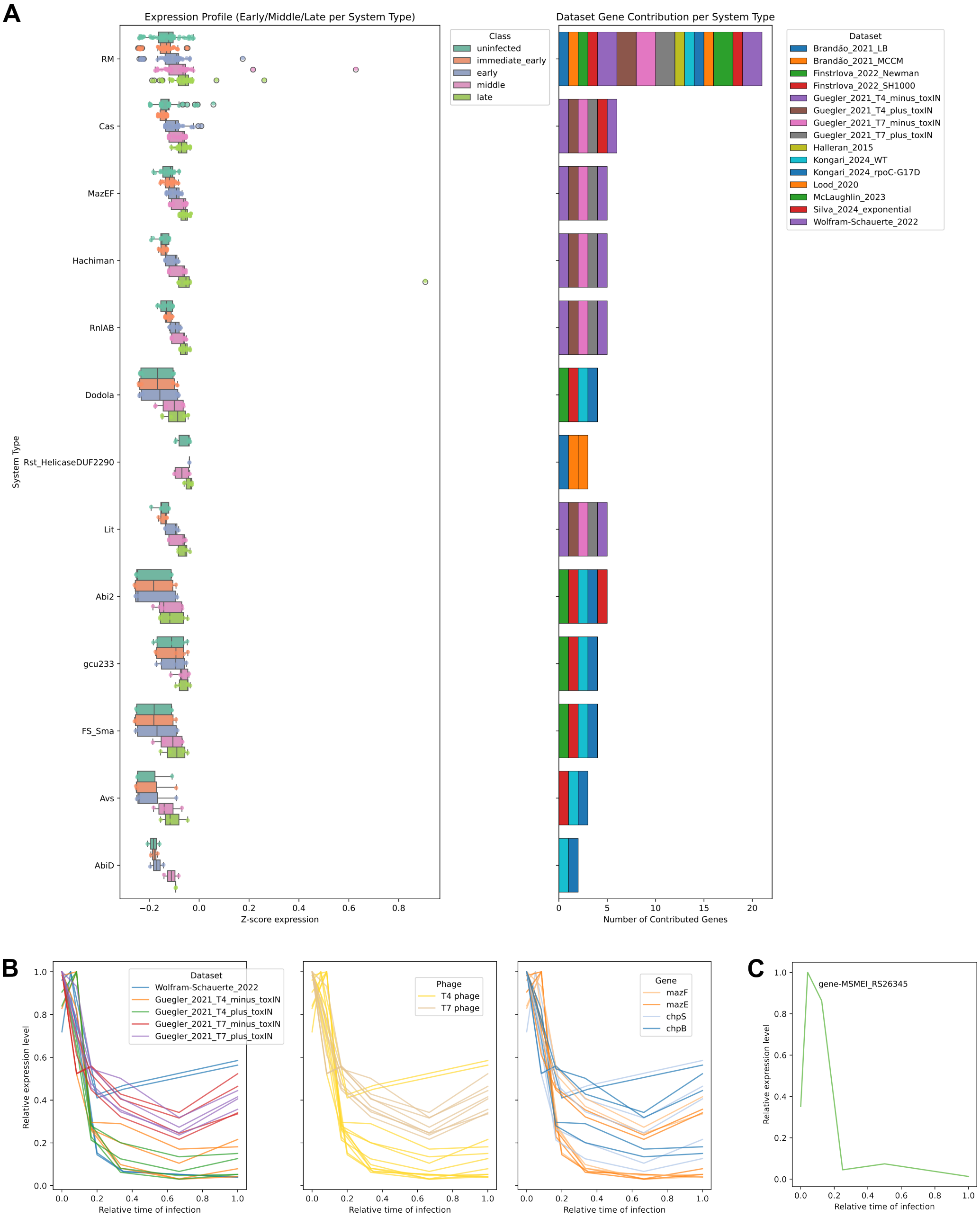


Supplementary Figure S13: **Host defense system expression.**
**A**) Boxplot of z-score normalized expression per sample of defense genes in uninfected hosts as well as in immediate early, early, middle and late infection phase (left). Stacked bar plot (right) indicates the number of genes contributed to each orthogroup by the individual datasets. **B**) Profile plot indicating the relative expression of the MazEF defense system in E. coli color by dataset (left), phage (middle) and gene (right). **C**) Profile plot of the relative expression of the single-gene defense system Ceres in *M. smegmatis*.

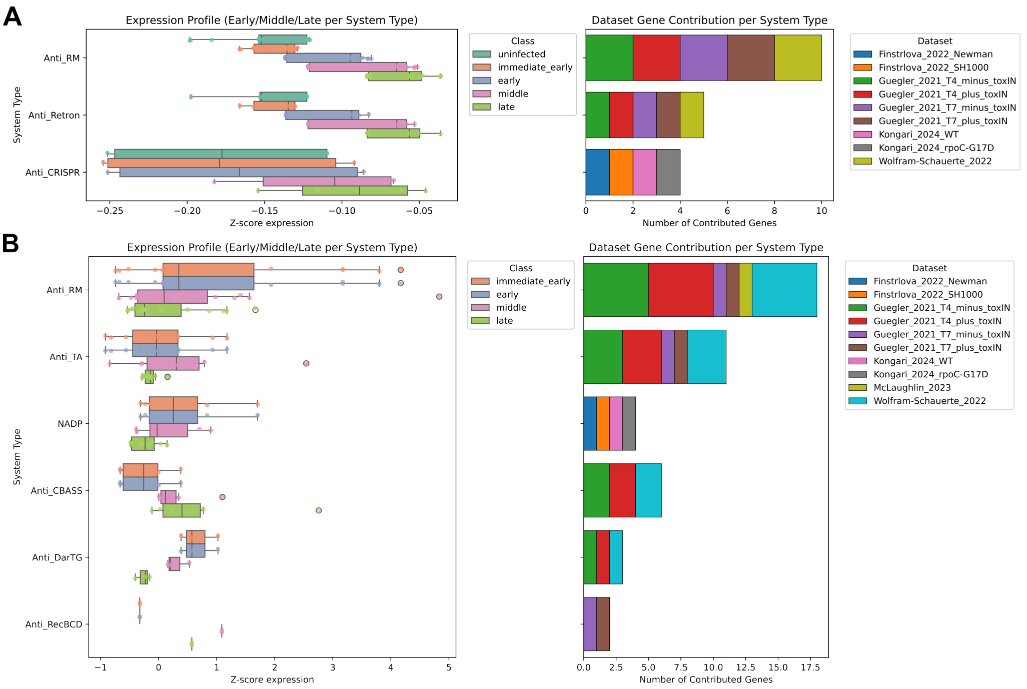


Supplementary Figure S14: **Host and phage anti-defense system expression.**
**A**) Boxplot of z-score normalized expression per sample of anti-defense genes in uninfected hosts as well as in immediate early, early, middle and late infection phase (left). Stacked bar plot (right) indicates the number of genes contributed to each orthogroup by the individual datasets. **B**) Same plot as in A for phage-encoded anti-defense systems.

### 2 Supplementary Tables

Supplementary Tables are available via Supplementary_Table_S1.xlsx.

- **Supplementary Table S1A**: Details on data collection for the PhageExpressionAtlas.
- **Supplementary Table S1B**: Details on dataset usage for phage gene classification benchmark and cross-host and -phage analysis.

### References


[1] M. Wolfram-Schauerte, N. Pozhydaieva, M. Viering, T. Glatter, and K. Höfer. Integrated omics reveal time-resolved insights into t4 phage infection of e. coli on proteome and transcriptome levels. *Viruses*, 14(11), 2022.

[2] A. Finstrlova, I. Maslanova, B. G. Blasdel Reuter, J. Doskar, F. Gotz, and R. Pantucek. Global transcriptomic analysis of bacteriophage-host interactions between a kayvirus therapeutic phage and staphylococcus aureus. *Microbiol Spectr*, 10(3):e0012322, 2022.
